# Relationship Between Gene Expression and Drug Response in Triple-Negative Breast Cancer: Leveraging Single-Cell RNA Sequencing and Machine Learning to Identify Biomarker Profiles

**DOI:** 10.64898/2026.03.05.709737

**Authors:** Kazhaleh Mohammadi, Nasim Afhami, Arthur Saniotis, Maciej Henneberg, Meisam Bagheri, Kaveh Kavousi

## Abstract

Triple-negative breast cancer (TNBC) is an aggressive subtype characterized by limited therapeutic options and poor prognosis. To address these challenges, we combined single-cell RNA sequencing (scRNA-seq) data with advanced machine learning techniques to find biomarkers that predict treatment response. Using tumor and blood samples from TNBC patients treated with either paclitaxel alone or in combination with atezolizumab, we constructed cellular maps, identified differentially expressed genes, and analyzed co-expression networks. Copy number variation (CNV) profiling and weighted gene co-expression network analysis (WGCNA) identified immune-related genes, including *IL7R*, *CD6*, and *TNFAIP3*, as candidate biomarkers linked to therapeutic response. A feature selection using Random Forest followed by bootstrap-enhanced K-nearest neighbors (K-NN) classification achieved high predictive accuracy across all subgroups (AUC > 0.93). Local Interpretable Model-Agnostic Explanations (LIME) based interpretability further identified important factors influencing treatment sensitivity and resistance, including *EGR1*, *MKI67*, *C1QA/B/C*, *GZMB*, and *PRF1*. Notably, blood-derived biomarkers showed strong predictive potential, highlighting the variability of liquid biopsy approaches for non-invasive monitoring. Our results indicate that integration of scRNA-seq with interpretable machine learning facilitates the identification of reliable biomarkers and yields clinically actionable insights for personalized therapeutic strategies in TNBC.

## 1. Introduction

Breast cancer remains the most prevalent malignancy among women globally and a leading cause of cancer-related mortality, primarily due to metastatic dissemination (Xu, 2021; Cai et al., 2025). Despite notable advancements in early detection and diversification of therapeutic options, a significant clinical challenge persists: the difficulty in accurately forecasting treatment outcomes at the individual patient level (Jain et al., 2024; Cummings-John et al., 2025). This challenge is most evident in aggressive molecular subtypes, where narrow therapeutic windows require highly precise, individualized prognostic tools that are often not provided by current clinical frameworks.

Standardized prognostic models frequently fail due to the inherent complexity of tumor heterogeneity. Breast tumors are not uniform masses; instead, they are dynamic ecosystems composed of diverse cellular subpopulations (Baba & Câtoi, 2007). This diversity leads to al heterogeneity, causing different clinical outcomes among patients, and intratumor heterogeneity, which is the presence of different lineages within a single tumor (Turashvili et al., 2017; Livesey & Marsh, 2022). From an evolutionary perspective, intratumor heterogeneity is influenced by selective pressures like nutrient limitation and immune surveillance. These pressures create "fitness tradeoffs" that enable certain clones to prosper (Hausser et al., 2019; Ahmed & Kim, 2024). Importantly, this heterogeneity often contains hidden, therapy-resistant subpopulations that endure initial treatments and lead to relapse (Garg et al., 2025). Therefore, any model ignoring this cellular "individuality" will lack the necessary resolution for effective precision medicine.

Traditionally, risk stratification models have relied on bulk RNA sequencing (RNA-seq). Although these methods have enhanced our understanding of overall transcriptomic patterns, their predictive power is limited by a "signal-averaging" effect (Wang et al., 2024). By averaging expression levels across millions of cells, bulk RNA-seq masks rare but clinically crucial cell states, such as cancer stem cells or chemo-resistant niches, which influence therapy outcomes. This reduced resolution hampers the ability to derive actionable mechanistic insights, often resulting in models that are statistically significant but lack biological clarity.

Single-cell RNA sequencing (scRNA-seq) has become a revolutionary approach to bridging this resolution gap. It measures the transcriptome at the single-cell level, enabling detailed deconstruction of the tumor microenvironment and uncovering specific gene regulatory networks that contribute to resistance, such as in triple-negative breast cancer (Zhang et al., 2021; Sun et al., 2021). Nevertheless, transforming this high-resolution data into clinical applications is challenging due to the high dimensionality and inherent noise in scRNA-seq datasets.

To translate this cellular granularity into predictive power, robust machine learning (ML) frameworks are required. ML algorithms are uniquely equipped to identify non-linear, multivariate relationships in large datasets, enabling the discovery of complex biomarkers that univariate analyses overlook (Cruz et al., 2007). By integrating the biological depth of scRNA-seq with the pattern-recognition capabilities of ML, we can move beyond descriptive observations and toward predictive models that accurately reflect the heterogeneous reality of breast cancer.

This study seeks to bridge the gap between high-resolution transcriptomics and clinical decision-making by identifying robust gene biomarkers that dictate drug response in Triple-Negative Breast Cancer (TNBC). By integrating single-cell RNA sequencing (scRNA-seq) with machine learning, we move beyond bulk-tissue averages to resolve how tumor heterogeneity and evolving cell states drive therapy resistance. Our approach enables the identification of distinct malignant subgroups with divergent sensitivities, offering a granular view of biomarker expression across various subtypes and cell cycle phases. By analyzing patient-derived expression profiles before and after drug exposure, we aim to define a predictive gene panel that guides personalized therapeutic interventions, ultimately ensuring that the right treatment reaches the right patient at the right stage of disease progression (Gambardella et al., 2022)."

## 2. Materials and Methods

### Single-Cell RNA Sequencing Data Preprocess

To investigate the relationship between gene expression and treatment response in triple-negative breast cancer (TNBC), a hybrid single-cell RNA sequencing analysis pipeline, including data preprocessing, batch fusion, and interpretive machine learning, was developed. This pipeline will identify key transcriptional biomarkers to predict treatment sensitivity or resistance. We obtained scRNA-seq GSE169246 (Zhang, Y., et al., 2021) and GSE161529 (Pal, B., et al., 2021) from the Gene Expression Omnibus (GEO) database (https://www.ncbi.nlm.nih.gov/geo/), Table 1. Specifically, the GSE169246 dataset, comprising 22 pairs of TNBC tumors and peripheral blood from advanced TNBC patients treated with paclitaxel or in combination with atezolizumab, was analyzed using the GPL20795 platform (HiSeq X Ten (Homo sapiens)). In addition, six normal epithelial samples from healthy donors (from GSE161529) were included as reference controls for copy number variation (CNV) analysis. CNV analysis was used to assess genomic changes in tumor tissue with treatment. The R packages “ggplot2” and “pheatmap” were used to visualize DEGs as volcano plots and heat maps.

**Table 1.**

| **Data source** | **Samples** | **Tissue** | **Platforms** | **References** |
| --- | --- | --- | --- | --- |
| **GSE169246** | **22** | **PBMCs / Tumor** | **10x Genomics** | **(Zhang, 2021)** |
| **GSE161529** | **6** | **Tumor** | **10x Genomics** | **(Chen, 2022)** |

**TNBC: Triple-Negative Breast Cancer**

### Single-cell RNA-sequencing dataset quality control and processing

Seurat v4.3.0 (Hao, Y., et al., 2021) in R was used for standard preprocessing and quality control. Cells with fewer than 500 identified genes or fewer than 2000 UMI counts were eliminated. Cells with more than 10% mitochondrial genes were also rejected. The goal of that selection was to eliminate low-quality cells. Next, SCTransform (Hafemeister, 2019) was used to stabilize variance and eliminate technical noise by normalizing counts using regularized negative binomial regression. Subsequently, Seurat’s anchor-based methods were employed to integrate data across treatment groups. By choosing the 3000 most variable genes for anchor identification and linking the datasets, it effectively corrected technical differences while preserving biological signals. Later, Principal Component Analysis (PCA) was applied for dimensionality reduction, with the top 40 principal components retained for further analysis. UMAP was then applied to these components to visualize cellular heterogeneity and to analyze clustering patterns and major cell types between pre- and post-treatment samples based on their transcriptomic profiles (McInnes, L. et al., 2018). Cell type annotation relied on the expression of standard marker genes and comparisons with external references such as Azimuth (Hao, Y., et al., 2021) and CellMarker (Zhang, X., et al., 2019). Importantly, for tumor data, the “inferCNV” R package was used to identify malignant cells with large-scale clonal chromosomal copy number variations (CNVs) (Patel, A. P., et al., 2014). Finally, the “intersect” function in R was used to identify common genes by overlapping the DEGs, module genes, and marker genes.

### Clustering and Differential Expression Analysis

To characterize transcriptional responses across cell types under different treatment conditions, we performed unsupervised clustering followed by cluster-level marker discovery. Cells were embedded in a shared nearest neighbor (SNN) graph constructed from principal component (PC) space, and clusters were identified using Seurat’s FindClusters () with a resolution of 0.8. A two-dimensional visualization was generated using UMAP.

In this analysis, cluster identities were assigned by comparison to reference atlases (Azimuth and CellMarker) and by expression of canonical marker genes, followed by manual expert review. Differential expression analysis was used to identify marker genes for each cluster using the Wilcoxon rank-sum test (Wilcoxon, 1946). This non-parametric test evaluates whether expression values for a given gene are systematically shifted between groups by comparing ranked observations. Marker genes were selected based on statistical significance and effect-size criteria, using thresholds on adjusted changes across relevant factors and p-value cutoffs. Gene annotations, including biological class and functional descriptions, were retrieved from Ensembl genomes (Cunningham, 2022). For each cluster, the top 10 markers were retained, and gene lists were exported as CSV files for downstream pathway enrichment and biomarker prioritization.

### Weighted Gene Co-expression Network Analysis

Weighted gene co-expression network analysis (WGCNA) was used as an initial strategy for biomarker prioritization (Langfelder, 2008). Co-expression modules were identified using the R package WGCNA, with minModuleSize = 30 and mergeCutHeight = 0.25.

To facilitate sample-level network architecture from scRNA-seq data, we derived pseudo-bulk expression profiles by aggregating transcriptomic signatures across individual cells within each biological sample. This approach preserves the sample-specific variance necessary for robust systems-level analysis while mitigating the technical noise inherent in single-cell datasets. Utilizing these profiles, we constructed independent gene co-expression networks for pre-treatment and post-treatment cohorts. By generating discrete networks for each clinical condition, we enabled the identification of condition-specific modules and the systematic comparison of topological shifts. This comparative framework allows us to resolve how drug intervention reconfigures gene regulatory programs and identify hub genes that drive therapeutic resistance or sensitivity.

Network construction followed standard WGCNA procedures. Briefly, pairwise gene correlations were transformed into an adjacency matrix using a soft-thresholding power selected to approximate scale-free topology, a property commonly observed in biological networks (Barabasi, 1999). The soft-thresholding exponent was set to β = 5, yielding a scale-free topology fit of R² = 0.9, based on the joint assessment of scale independence and mean connectivity. The adjacency matrix was then converted into a Topological Overlap Matrix (TOM), which quantifies network interconnectedness by measuring the degree of shared neighbors between gene pairs (Nguyen, P., et al., 2025).

Genes were clustered hierarchically using TOM-based dissimilarity, and modules were defined using dynamic tree-cutting. To evaluate treatment-associated module changes, we computed Pearson correlations between module eigengenes from the pre-treatment network and the post-treatment network to identify corresponding and/or reconfigured modules across conditions. We then assessed associations between post-treatment module eigengenes and treatment response traits. Finally, for each pre-treatment module, we calculated a module importance score by integrating (i) its correspondence to post-treatment modules and (ii) the strength of trait association observed in the post-treatment network (Horvath, 2011).

Candidate biomarkers were prioritized based on module membership (kME) within treatment-associated modules, with higher module membership indicating stronger connectivity to the module’s eigengene and greater relevance to the module’s co-expression program. Genes with high membership in trait-associated modules were retained for downstream machine learning analyses. All code used for pseudo-bulk generation, network construction, and downstream integration is publicly available at: https://github.com/KazhalehMohammadi/Tumor_TNBC_scRNAseq/.

### Functional Enrichment Analysis

Functional enrichment analysis was performed using the Gene Ontology (GO) (Consortium, 2021) and Kyoto Encyclopedia of Genes and Genomes (KEGG) (Kanehisa, M., et al., 2021) databases to evaluate the biological significance of DEGs for each condition separately. The DEGs were grouped into four conditions: tumor paclitaxel, tumor paclitaxel & atezolizumab, blood paclitaxel, and blood paclitaxel & atezolizumab. Enrichment analysis was performed using pre-computed DEG lists for each condition with the cluster Profiler (Yu, G., et al., 2012) and “org.Hs.eg.db” (Huber, W., et al., 2015) packages in R, filtering results to include only pathways with adjusted P-values < 0.05.

To delineate the functional landscape of the treatment response, we performed a comparative enrichment analysis of Gene Ontology (GO) and KEGG pathways across both datasets. This approach allowed us to resolve the convergence and divergence of signaling programs between the two experimental conditions. Shared and condition-specific pathways were rigorously validated using set-theoretic operations, specifically intersection and set-difference analysis, to isolate conserved biological drivers from unique treatment-induced effects. The statistical significance and overlap of these terms were prioritized using clustered heatmaps of -log_10_ (adjusted p-values), providing a granular view of pathway activity. To highlight the most biologically relevant transitions, the top 10 significantly enriched GO and KEGG pathways within the treatment cohort were visualized via dot plots, emphasizing the magnitude of enrichment and the associated gene ratios.

An integrated summary table was created by compiling data from the most frequently enriched pathways with the highest significance, based on their frequency and lowest adjusted p-value. This table presents the top 10 GO and KEGG pathways for each group, facilitating comparisons of the biological functions affected by paclitaxel alone versus combination therapy.

### Biomarker Prioritization

Biomarker prioritization was performed using a predefined multi-layer scoring framework that integrated multiple analytical outputs. Candidate genes were selected based on evidence scores, and WGCNA module membership was a major additional consideration when many genes showed similarly patterned expression. WGCNA identifies genes associated with treatment-related co-expression modules. Biomarkers in TNBC were identified using this integrative strategy, which combined scores from differential gene expression, weighted gene co-expression network analysis, pathway enrichment, and high-confidence CNV analyses. This strategy was applied across two treatment arms: tumor-paclitaxel and tumor-paclitaxel & atezolizumab. To further strengthen prioritization, the biological support for each gene was quantified by assigning an evidence score. This score was calculated as the sum of the following components:

Evidence Score = (Significant in GO Enrichment) + (Significant in KEGG Enrichment) + (Located in CNV-altered region (Tumor data)) + (Significant in DEG)

We assigned one point to each gene for every condition it met. Genes with higher scores were considered more biologically relevant, supported by multiple independent lines of evidence. For each group, the top 100 biomarkers were ranked according to these scores. In case of ties, the WGCNA module membership (kME) was used as a tiebreaker, favoring genes more strongly linked to treatment-related co-expression networks.

## Machine Learning Techniques

### Machine learning-based screening of candidate diagnostic genes

We developed a two-step machine learning process to identify biomarkers predicting therapeutic response across four treatment subgroups: tumor–paclitaxel, tumor–paclitaxel & atezolizumab, blood–paclitaxel, and blood–paclitaxel & atezolizumab. scRNA-seq data from TNBC patients in each group, as mentioned in the previous section on Single-cell RNA-sequencing dataset quality control and processing, were thoroughly quality-checked and preprocessed. Subsequently, a Random Forest algorithm was used to select the most informative genes. The Random Forest method was chosen because it is a robust ensemble classifier that combines predictions from multiple decision trees built from randomly selected gene subsets. Each tree independently predicts a class, and the overall output is determined by majority voting among all trees (Moorthy, K., et al., 2011).

Mathematically, the Random Forest classifier predicts the class label based on the output of T trees as follows:

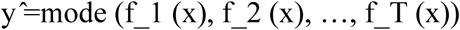

Where f_t (x) denotes the class predicted by tree t for sample x, feature importance scores were calculated as the overall reduction in node impurity attributed to each feature across the forest, thus allowing genes to be ranked according to their contribution to treatment-response discrimination.

For prediction evaluation, bootstrapping was used. For each bootstrap iteration, variable importance scores were estimated on the training data, and the 20 most significant genes were selected. At the end of all iterations, the most frequently selected genes were reported as candidate diagnostic biomarkers. This process reduced data complexity while focusing on genes most linked to treatment outcomes (Efron, 1997) (Díaz-Uriarte, 2006).

### Bootstrap and K-NN Classification

We implemented a bootstrap-based classification framework using K-Nearest Neighbors (K-NN) to assess the predictive value of the selected biomarkers. Each treatment subgroup was analyzed independently across 100 bootstrap iterations. Each iteration involved selecting a training set with 70% of the observations and a testing set with the remaining 30%. Each bootstrapped iteration followed a systematic modelling and assessment methodology.

To ensure the stability of our predictive gene panel, we implemented a multi-stage feature selection pipeline integrated within a bootstrapping framework. For each iteration, a Random Forest (RF) regressor was employed to rank genes based on their mean decrease in impurity (Gini importance) or permutation importance. From this ranked distribution, the top 20 most salient features were prioritized and utilized to train a K-Nearest Neighbors (K-NN) classifier. The K-NN algorithm then predicted class labels using a majority-vote heuristic across the k nearest neighbors in the high-dimensional feature space. By coupling the non-linear feature ranking of Random Forest with the local-topology-based classification of K-NN across multiple bootstrap resamples, we minimized the risk of overfitting and ensured the generalizability of our treatment-response signatures.

To determine the class label y ^ for a given sample x, the K-NN classifier utilizes a majority-vote heuristic defined as:

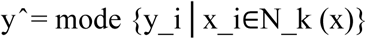

where N_k(x) denotes the set of k-nearest neighbors in the feature space. To mitigate biases inherent in imbalanced datasets, particularly the disproportionate representation of drug-resistant versus sensitive samples, we integrated the Synthetic Minority Over-sampling Technique (SMOTE) into the training phase. Rather than simple duplication, SMOTE synthesizes new instances by interpolating between minority-class observations and their nearest neighbors (Chawla et al., 2002). This synthetic augmentation helps ensure a more equitable decision boundary. Finally, to ensure the model’s generalizability, hyperparameters were tuned using a 5-fold cross-validation strategy, optimizing for the most stable predictive performance.

Model performance was evaluated using bootstrap test sets. Metrics assessed included accuracy, sensitivity, specificity, precision, F1-score, and the area under the ROC curve (AUC). Results were averaged over 100 iterations to obtain stable estimates of predictive performance for each subgroup. The integrated bootstrap-KNN framework enabled simultaneous feature selection and classification, thereby minimizing bias and providing robust, reliable estimates of treatment-specific biomarkers.

### Interpreting and selecting the top genes using LIME

To improve the transparency of the machine learning model and provide better insights into its predictions, the Local Interpretable Model-agnostic Explanations (LIME) framework was applied. LIME approximates the behavior of a complex model by fitting simple, interpretable models to each individual observation. Therefore, LIME can provide a clear view of how each gene contributes to the prediction outcome for specific samples, allowing us to better understand the model’s decision-making process.

To facilitate interpretation, LIME was applied to the K-NN classifier using only the top 20 most selected genes from the feature selection output in the previous step. By focusing the analysis on these 20 biomarkers, the model’s interpretability was enhanced by highlighting biologically relevant features. LIME was applied to randomly selected test samples to illustrate the relative influence of each gene on classification. The following descriptions identified genes with the strongest positive or negative effects on predictions, enabling us to rank a subset of candidate biomarkers. Based on these analyses, the most consistently significant genes were highlighted as candidate biomarkers associated with therapeutic response in TNBC with different treatment regimens.

### Statistical analysis

All data processing and statistical analyses were performed using R software (version 4.3.0). Differential gene expression analysis between groups was conducted using the Wilcoxon rank-sum test, as implemented in the Seurat package. P-values were adjusted for multiple testing using the Bonferroni correction unless otherwise stated. For correlation-based analyses, including weighted gene co-expression network analysis (WGCNA), Pearson correlation coefficients were calculated. Quantitative results are reported as mean ± standard deviation (SD) unless otherwise specified. The statistical significance was defined as a p-value < 0.05.

## 3. Results

### Characterization of cellular composition changes across tumor and peripheral blood under paclitaxel and paclitaxel plus atezolizumab

To resolve the cellular architecture of triple-negative breast cancer (TNBC) under therapeutic pressure, we used single-cell RNA sequencing (scRNA-seq) data from 388,049 high-quality cells. These cells were captured from both tumor and peripheral blood mononuclear cell (PBMC) samples across two distinct cohorts: paclitaxel monotherapy and paclitaxel combined with atezolizumab (anti-PD-L1). High-dimensional visualization via UMAP revealed profound cellular heterogeneity, categorized into 106 transcriptional subclusters representing 12 primary lineages identified by canonical marker expression. The cellular subclusters were identified using a combination of automated label transfer and manual marker-based rules, ensuring accurate annotation of cell types and lineages. The workflow chart is presented in Figure 1.

**Figure 1:**
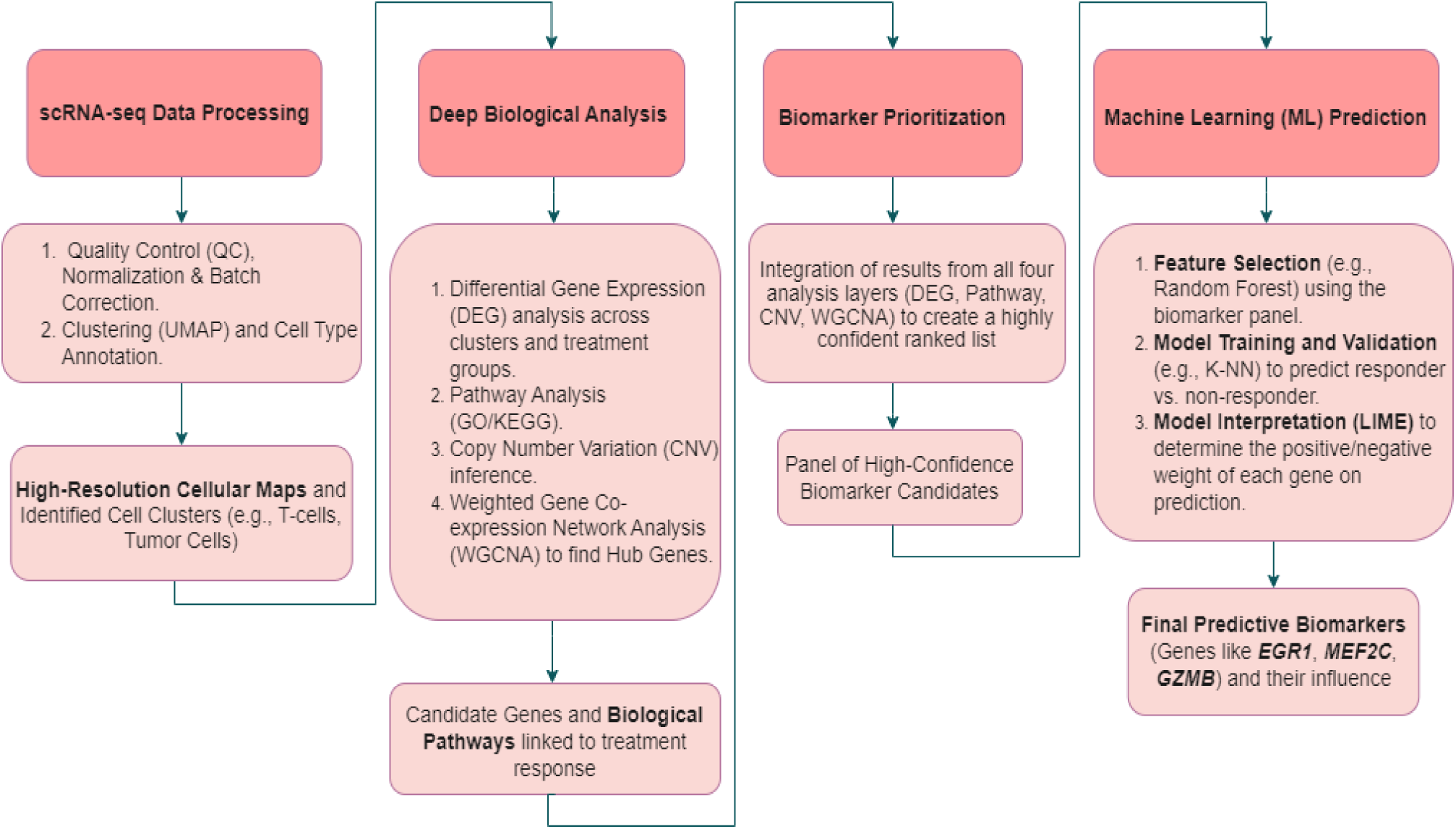
High-Definition Workflow for Predictive Biomarker Discovery in Individualized Treatment Response using scRNA-seq Data. The process here began with scRNA-seq Data Processing, that is, quality control, normalization, clustering, and annotation of cell types into high-resolution cellular maps that define cell clusters. The subsequent steps involved Deep Biological Analysis, including differential gene expression analysis, pathway analysis, copy number variation inference, and weighted gene co-expression network analysis to identify hub genes. These biological insights were integrated into Biomarker Prioritization, leading to the selection of a high-confidence biomarker candidate panel. Finally, Machine Learning Prediction was employed to perform feature selection, model training and validation, and model interpretation, ultimately identifying predictive biomarkers and assessing their impact on treatment response.

### UMAP-Based Characterization of Tumor Cell Composition Under Paclitaxel Monotherapy and Paclitaxel Combined with Atezolizumab

UMAP analysis of tumor samples from patients treated with paclitaxel monotherapy identified 10 transcriptionally distinct cell populations, including malignant epithelial cells, fibroblasts, activated T cells, cytotoxic T cells, exhausted T cells, regulatory T cells, NK cells, B cells, macrophages, and dendritic cells (Figure 2A). These populations formed well-separated clusters, indicating transcriptional differences among immune, stromal, and malignant compartments.

**Figure 2:**
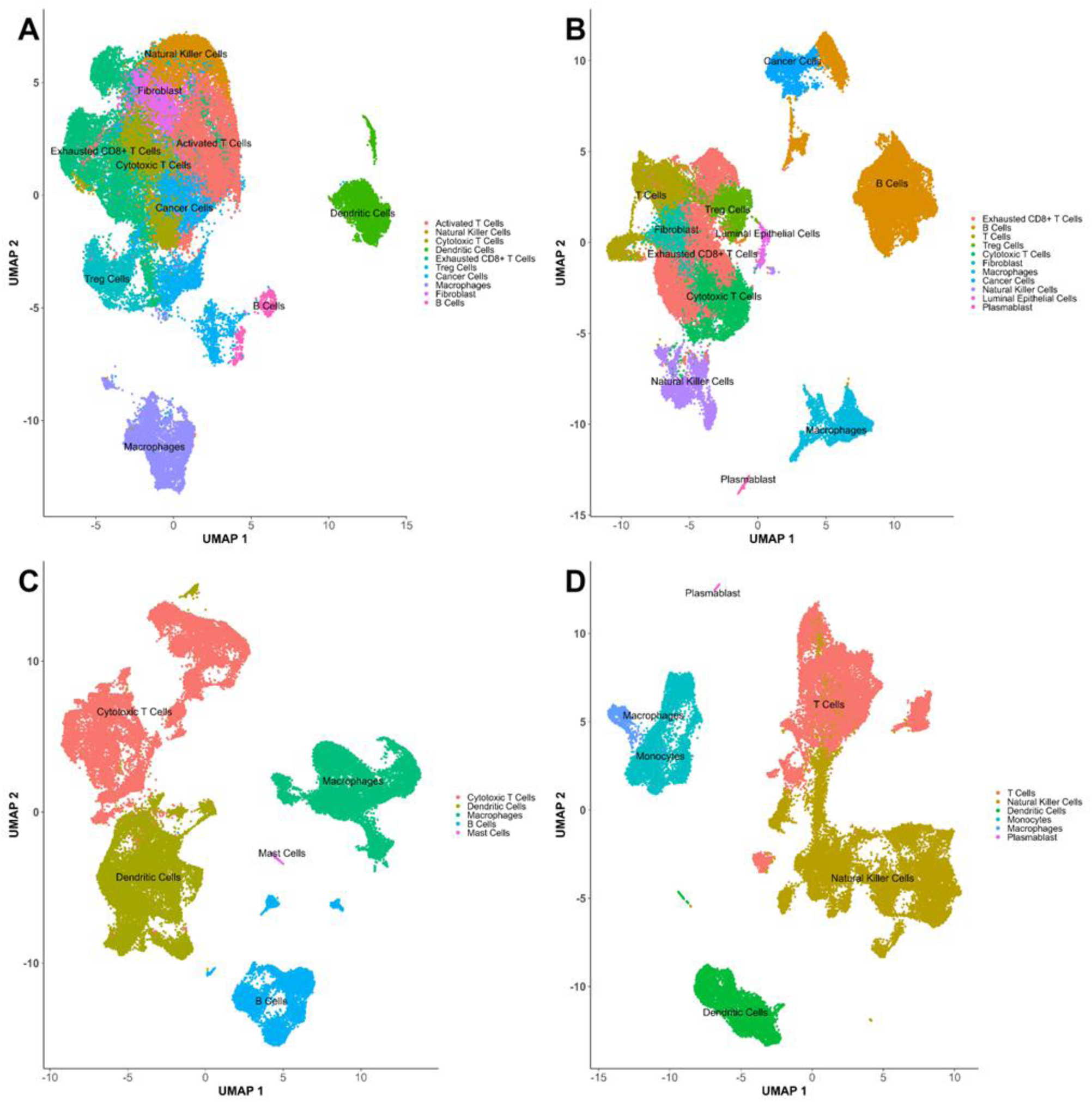
**(A)** UMAP representation of various cell populations in tumor samples from patients treated with only paclitaxel. Results show the distribution of different cell types, including Activated T Cells, Cytotoxic T Cells, and Cancer Cells. **(B)** Tumor cell populations in UMAP representation from patients treated with paclitaxel and atezolizumab combination therapies. Significant alteration in cell type distribution was noted when compared to paclitaxel-only treatment. **(C)** UMAP analysis of blood samples from patients treated with paclitaxel, showing the distribution of various blood cell types, including cytotoxic T cells and B cells. **(D)** UMAP analysis of blood samples from those treated with a combination of paclitaxel and atezolizumab. Compared to the paclitaxel-only treatment, there are notable changes in the distribution of the blood cell populations.

In tumor samples subjected to combination therapy (paclitaxel and atezolizumab), the fundamental lineage architecture remained conserved; however, significant shifts in the relative abundance of specific cell populations emerged (Figure 2B). Comparison of percentage changes in gene expression between paclitaxel monotherapy and combination treatment revealed a significant effect exclusively within the macrophage compartment (mean change = +21.1%, p = 0.035, Wilcoxon rank-sum test). In contrast, no statistically significant differences were detected for B cells, cytotoxic T cells, or total T cells (p > 0.05 for all comparisons), indicating that the addition of atezolizumab did not substantially alter transcriptional dynamics within these lineages. Differential expression patterns in macrophages under combination therapy suggested a treatment-specific modulation of innate immune responses, while other lymphoid populations remained transcriptionally comparable to those observed under paclitaxel alone. Collectively, these findings indicate that although combination therapy preserves global cellular diversity, its principal impact on the tumor microenvironment is mediated through selective reprogramming of macrophage-associated gene expression rather than broad lymphoid expansion (Table 2).

**Table 2.**

| Group | Mean_Change_Tumor | P_Value_Tumor |
| --- | --- | --- |
| B Cells | 0.8854736 | 0.6305289 |
| Macrophages | 21.1471339 | 0.0354630 |
| Cytotoxic T Cells | 7.0130568 | 0.6305289 |
| T Cells | -2.7809754 | 0.8315005 |

Tumor Cell Composition Under Paclitaxel Monotherapy and Paclitaxel Combined with Atezolizumab

### UMAP-Based Characterization of Peripheral Blood Immune Cell Composition Under Paclitaxel Monotherapy and Paclitaxel Plus Atezolizumab

To evaluate whether the local tumor microenvironment (TME) remodeling was mirrored by systemic immune shifts, we profiled the circulating immune landscape via peripheral blood analysis. Under paclitaxel monotherapy, the systemic compartment maintained a stable architecture across five primary lineages: cytotoxic T cells, monocyte-derived cells, B cells, mast cells, and dendritic cells (Figure 2C). Conversely, combination therapy triggered a distinct systemic reconfiguration, characterized by NK cells, monocytes, and plasmablasts (Figure 2D).

In blood samples treated with paclitaxel and atezolizumab, the overall cellular architecture remained largely consistent, but notable changes in specific cell populations were observed. Analysis of percentage changes in gene expression between paclitaxel monotherapy and combination treatment revealed a significant reduction in macrophage populations (mean change = -21.24%, p = 1.3e-15, Wilcoxon rank-sum test). Conversely, no significant changes were observed in dendritic cells (mean change = -0.12%, p > 0.05), suggesting that the addition of atezolizumab had minimal impact on this lineage. These findings highlight a selective decrease in macrophages following combination therapy, while dendritic cells and other lymphoid populations, including cytotoxic T cells, did not show substantial alterations. Overall, while the combination therapy did not broadly affect the immune landscape in blood samples, it led to a significant reduction in macrophage-related gene expression, implying that the therapeutic effect may be more closely tied to the modulation of macrophage activity rather than broad immune expansion.

Lineage fidelity and functional states were rigorously assigned using canonical markers, including IL7R, FOXP3, CD8A, GZMB, and HLA-DRA. To isolate high-confidence signatures, we employed the Wilcoxon rank-sum test with Bonferroni correction, enforcing stringent thresholds (p < 0.05; |log_2FC| > 1). While heatmap visualization confirmed that core lineage identities remained robust across cohorts, it revealed a clear divergence in marker expression intensity driven by the combination regimen (Figure 3). This systemic surge in plasmablasts aligns with our intra-tumoral observations, suggesting a coordinated B-lineage response to the chemo-immunotherapy dual block (Table 3).

**Figure 3:**
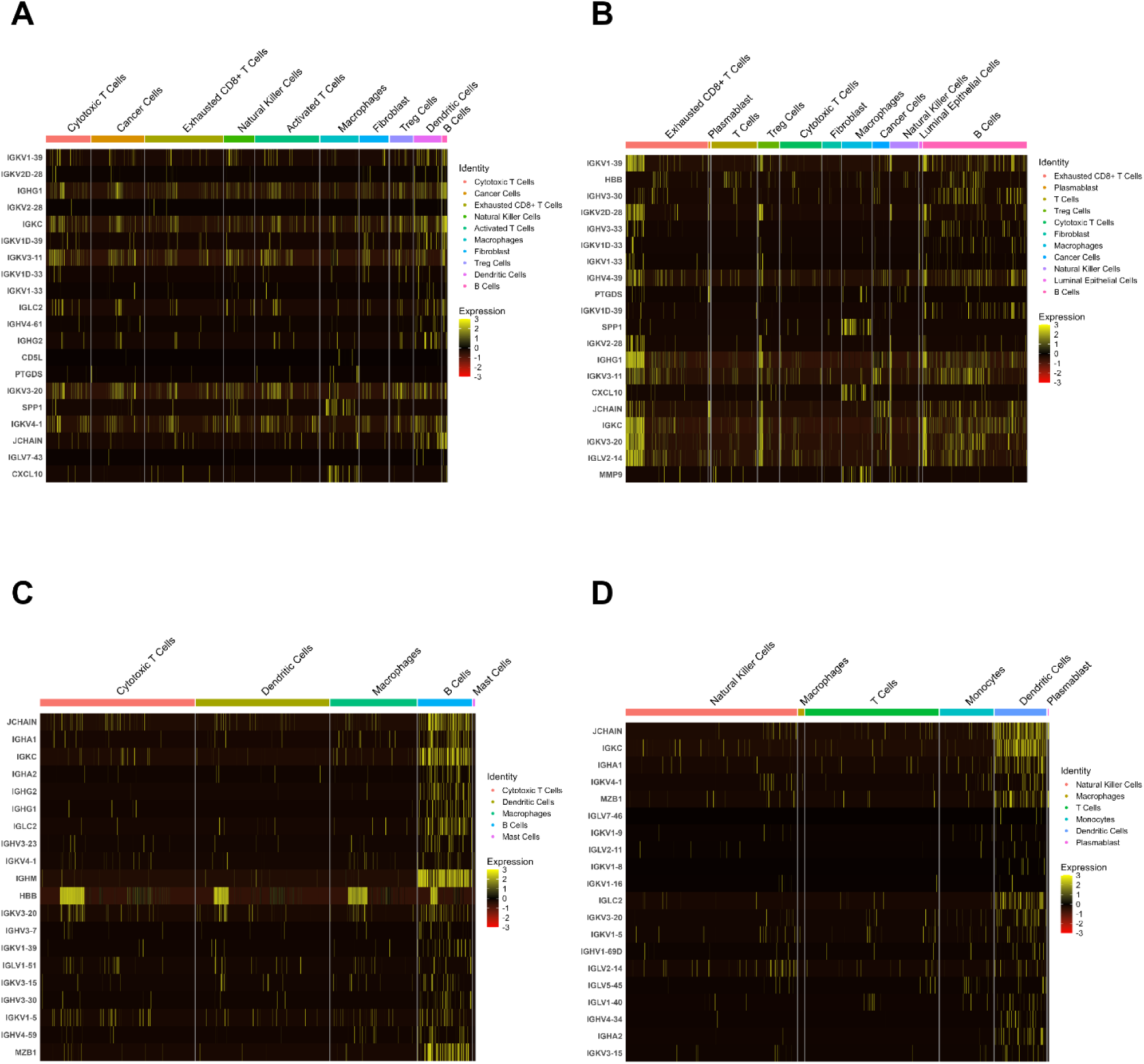
**(A)** Tumor sample under paclitaxel monotherapy treatment. **(B)** Tumor sample under paclitaxel combined with atezolizumab treatment. **(C)** Blood sample under paclitaxel monotherapy treatment. **(D)** Blood sample under treatment with paclitaxel combined with atezolizumab.

**Table 3.**

| Group | Mean_Change_Blood | P_Value_Blood |
| --- | --- | --- |
| Dendritic Cells | -0.1167011 | 0.4755354 |
| Macrophages | -21.2442503 | $1.30 \times 10^{-15}$ |
| Peripheral blood Cell Composition Under Paclitaxel Monotherapy and Paclitaxel Combined with Atezolizumab |  |  |

### Differential Transcriptional Landscapes and Functional Diversification

To delineate the treatment-specific transcriptional programs in TNBC, we performed comparative differential expression analysis across tumor and peripheral blood mononuclear cell (PBMC) compartments. Analysis of tumor samples revealed a distinct transcriptomic divergence between the two regimens (Supplementary Table 1). Notably, the chemokine XCL1—a key recruiter of cross-presenting dendritic cells—exhibited a contrasting trajectory: it was downregulated following paclitaxel monotherapy but significantly upregulated in the combination group (log_2FC difference = 1.46). Similarly, the B-lineage markers IGKV1-39, IGHG1, and IGKC showed robust positive shifts in the combination arm compared to monotherapy (differences of 1.40, 1.20, and 0.69, respectively), suggesting a synergistically induced B-cell response within the tumor niche. Systemic immune profiling of PBMC samples further mirrored this compartmentalization. Under paclitaxel monotherapy, significant modulation was restricted to a narrow set of inflammatory markers, including S100A9 and LYZ, while mast cell-associated transcripts (CPA3, HPGDS, TPSAB1) were consistently suppressed. In contrast, the combination therapy cohort displayed a more expansive systemic response. We observed substantial induction of IGHA1 (+0.86), PTGDS (+0.81), and the chemoattractant CCL4 (+0.69). Paradoxically, the pro-inflammatory alarmins S100A8 (-1.06) and S100A9 (-1.01), along with the early-response transcription factor FOS (-0.80), were sharply downregulated post-combination treatment, potentially indicating a resolution of systemic myeloid-driven inflammation. Functional enrichment analysis corroborated these gene-level shifts, revealing distinct pathway signatures across conditions (Supplementary Table 2). Paclitaxel monotherapy primarily enriched for baseline leukocyte-mediated immunity within the tumor (Figure 4A) and cell killing in the periphery (Figure 4C). However, the addition of atezolizumab fundamentally rewired these signatures. Tumors under combination therapy were characterized by "chemokine signaling" and cytokine-receptor interactions (Figure 4B), while the systemic landscape transitioned toward B-cell-mediated immunity and humoral immune responses (Figure 4D). KEGG analysis further confirmed this systemic shift, with significant enrichment in hematopoietic cell lineage and Th1/Th2 cell differentiation, suggesting that the combination regimen promotes a more sophisticated, multi-lineage immune mobilization compared to the cytotoxic effect of monotherapy alone.

**Figure 4:**
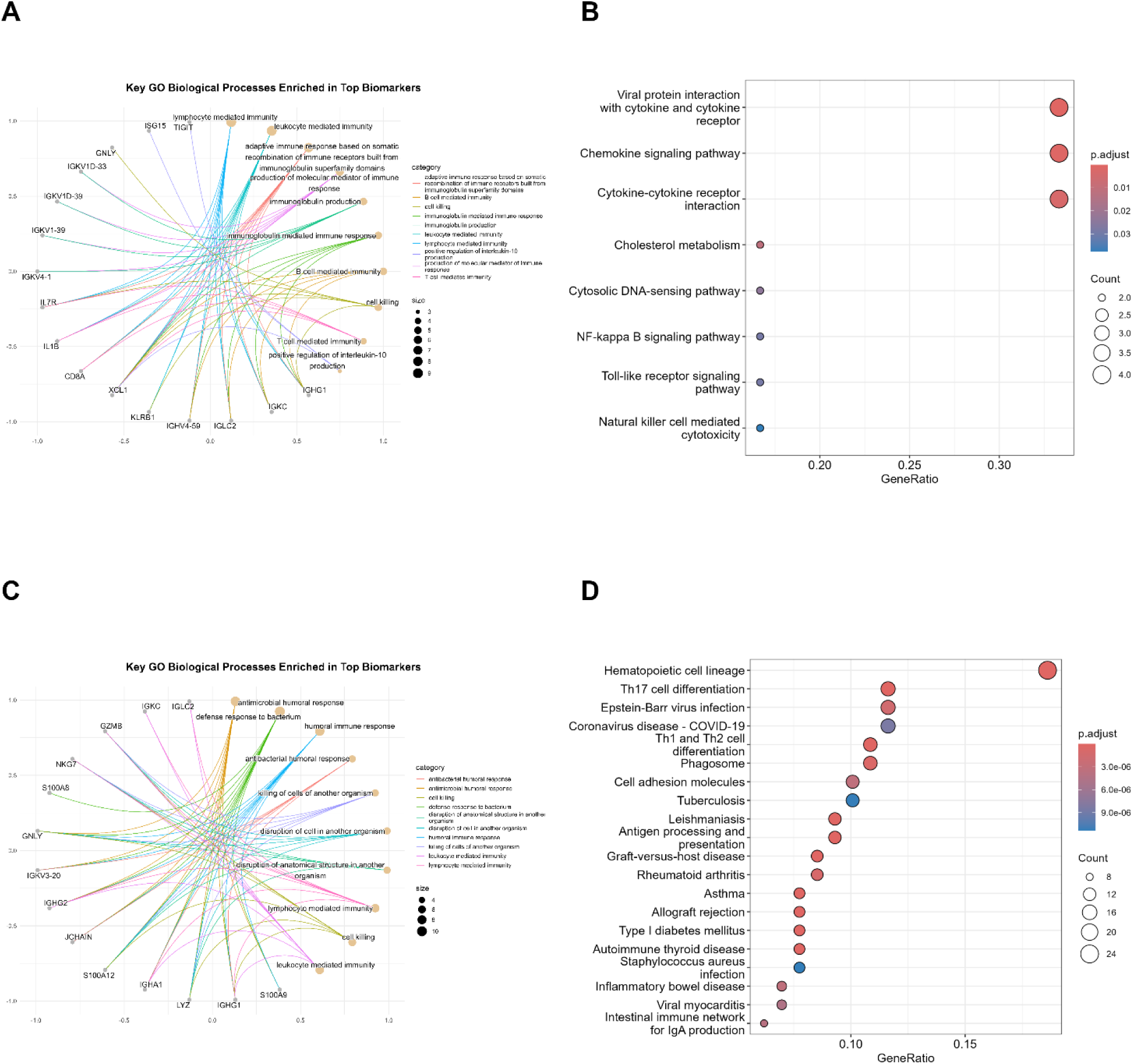
The image overall demonstrates the different biological and immune responses activated by treatment with paclitaxel and atezolizumab in triple-negative breast cancer patients. The results show that paclitaxel-only treatment primarily activates immune pathways such as lymphocyte-mediated immune response, while the combination of paclitaxel and atezolizumab activates more complex pathways like viral protein interactions with cytokines and chemokine signaling. Additionally, in blood samples, the combination therapy clearly activates pathways related to blood cell production and immune responses, highlighting the enhanced immune activation with combination therapy. **(A)** Cnetplot shows the significant Gene Ontology biological processes associated with the top biomarkers in tumor samples treated with only paclitaxel from triple negative breast cancer patients. As it is seen, different immune response pathways are shown in this plot, such as lymphocyte-mediated immune response and immunoglobulin-mediated immune response. **(B)** Dotplot showing KEGG pathways enriched in tumor samples from TNBC patients treated with the combination of paclitaxel and atezolizumab, highlighting key pathways such as viral protein interactions with cytokine and cytokine receptor, and chemokine signaling. **(C)** Cnetplot shows the key Gene Ontology biological processes enriched in top biomarkers in blood samples from TNBC patients treated with paclitaxel, including B cell-mediated immunity, cell killing, and humoral immune response. **(D)** KEGG pathways enriched in blood samples from triple negative breast cancer patients treated with the combination of paclitaxel and atezolizumab are shown through Dotplot. It is very obvious that the activation of pathways concerning blood cell production jobs like hematopoietic cell lineage, Th17 cell differentiation, and Th1/Th2 differentiation indicates serious participation of the immune system during disease progression.

### Genomic Architecture and Co-expression Network Dynamics

To investigate the genomic drivers of treatment response, we first performed single-cell CNV analysis. In paclitaxel-treated tumors, we identified significant expression alterations in genes localized to altered genomic regions, notably IGHG1, IGKC, and IGKV1-39 within B-cell lineages, and APOE and CST3 within the macrophage compartment (Figure 5A). These shifts suggest that paclitaxel-induced stress may selectively modulate the transcriptomic output of specific genomic loci. The addition of atezolizumab further accentuated these alterations. Immune-modulatory genes within macrophages showed the most profound shifts, with CST3 (FC = 5.39), APOC1 (FC = 5.53), and APOE (FC = 6.44) exhibiting substantial upregulation (Figure 5B). This dramatic induction of the APOE/APOC1/CST3 axis genes, frequently associated with lipid metabolism and immunosuppressive myeloid states, suggests a localized metabolic reprogramming of the TME. Pathway enrichment of these CNV-associated genes highlighted key roles in immune cell activation and inflammatory signaling (Supplementary Table 3), indicating that the combination regimen targets the genomic-transcriptomic interface to reshape treatment efficacy.

**Figure 5:**
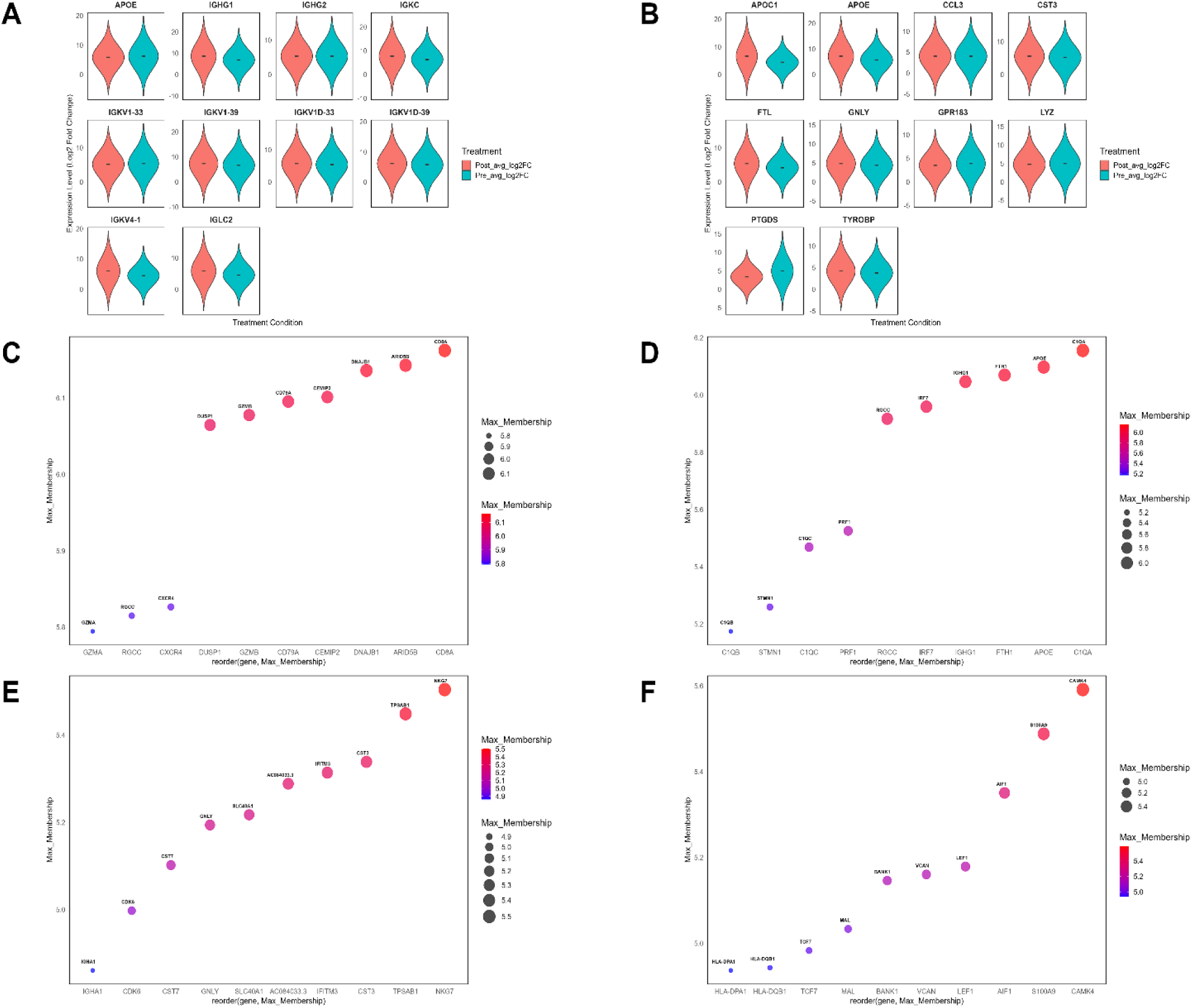
**(A)** CNV analysis reveals significant alterations in immune-related genes post-treatment with paclitaxel. Genes such as IGHG1, IGKC, and APOE were identified in B cells, showing substantial changes in expression, particularly in log2 fold change, indicating their role in immune modulation and inflammatory response to treatment. **(B)** CNV analysis reveals significant alterations in immune-related genes post-treatment with paclitaxel & Atezolizumab. Genes such as APOE, APOC1, and CST3 were identified in macrophages, showing substantial changes in expression, particularly in log2 fold change, indicating their role in immune modulation. **(C)** WGCNA analysis on tumor samples treated with paclitaxel identifies CD8A, ARID5B, DNAJB1, and CEMIP2 as hub genes within immune and cell signaling networks. These genes play pivotal roles in regulating immune response and cell migration. **(D)** WGCNA of tumor samples treated with paclitaxel and atezolizumab shows that genes such as DNAJB1, IL7R, TNFAIP3, and C1QA are crucial in regulating treatment response and disease progression. **(E)** In blood samples treated with paclitaxel only, GNLY, FGFBP2, and GZMA show significant changes, indicating their involvement in immune function and inflammation. **(F)** For blood samples treated with paclitaxel and atezolizumab, genes such as CAMK4, AIF1, and S100A9 demonstrated high module membership values, highlighting their roles in immune signaling, macrophage activation, and T-cell responses.

To resolve the higher-order organization of these genes, we utilized Weighted Gene Co-expression Network Analysis (WGCNA). In tumors treated with paclitaxel monotherapy, the co-expression landscape was dominated by modules centered on CD8A, GZMB, and CXCR4, reflecting a baseline cytotoxic immune signature (Figure 5C). However, combination therapy shifted the network topology toward modules anchored by DNAJB1, IL7R, and C1QA (Figure 5D). The emergence of C1QA as a hub gene is particularly salient, as it marks a transition toward a complement-mediated and potentially more robust anti-tumor response. Systemic network analysis of PBMC samples revealed a similar divergence. Paclitaxel monotherapy identified GNLY (MM = 5.19), GZMA (MM = 3.88), and FGFBP2 (MM = 3.49) as primary hubs, consistent with a circulating cytotoxic T-cell and NK-cell signature (Figure 5E). In contrast, the combination therapy group exhibited a significantly more complex circulating network. Hub genes in this cohort included CAMK4 (MM = 5.59), S100A9 (MM = 5.49), AIF1 (MM = 5.35), and BANK1 (MM = 5.15) (Figure 5F). The high connectivity of BANK1 (a B-cell scaffold protein) and CAMK4 (involved in T-cell signaling) suggests that the combination regimen facilitates a highly integrated, multi-lineage systemic response that is absent in monotherapy.

Using an integrated multi-level analytical framework, a set of candidate biomarkers with predictive relevance to treatment response in TNBC was identified. Biomarker prioritization was based on the convergence of multiple analytical layers, including differential gene expression, co-expression network membership, CNV-associated transcriptional changes, and functional pathway enrichment. The resulting panels comprised high-confidence biomarkers supported by at least 2 independent lines of evidence within each treatment and sampling group.

The genes such as *CD8A, CD79A*, and *GZMB,* in the paclitaxel treated tumor group, which are related to immune response and cellular toxicity, were identified. These genes are particularly involved in cytotoxic T cells, dendritic cells, and, Exhausted CD8+ T Cells, and are important markers of tumor immune activation and exhaustion (Figure 6A). While in the paclitaxel and atezolizumab treated tumor group, genes such as *IRF7, PRF1*, and *APOE*, which refer to the combination of immune activation and macrophage accumulation, were identified. These genes indicate the effect of the combination of paclitaxel and atezolizumab on enhancing immune responses and its effects on immune cells in the tumor (Figure 6B).

**Figure 6:**
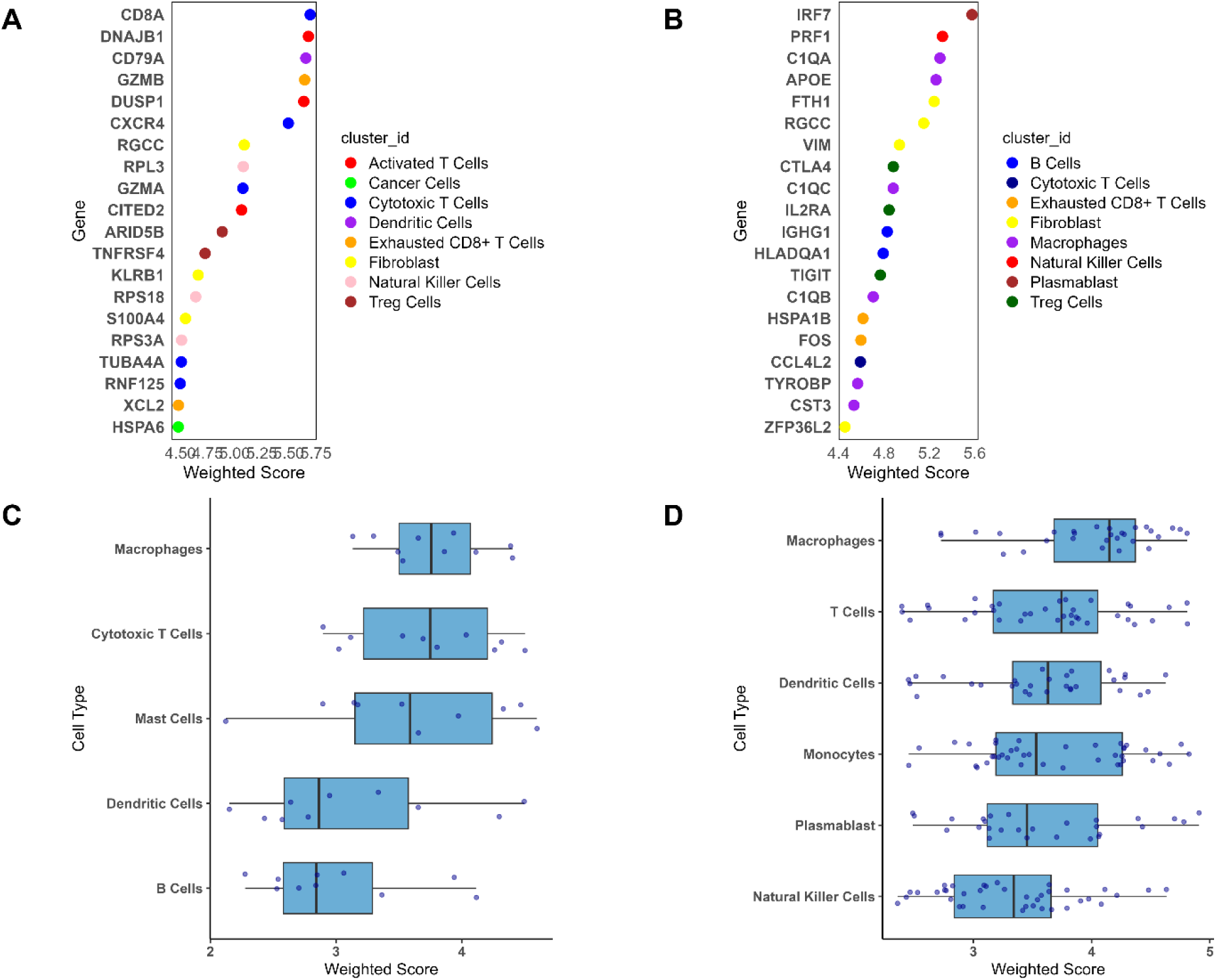
(A) Identification of key biomarkers in tumor and blood groups. **(A)** Key biomarkers in the paclitaxel-treated tumor group (CD8A, CD79A, GZMB) linked to immune response and toxicity, particularly in cytotoxic T cells and exhausted CD8+ T cells. **(B)** In the paclitaxel and atezolizumab-treated tumor group, IRF7, PRF1, and APOE indicate immune activation and macrophage accumulation, enhancing immune responses. **(C)** For the paclitaxel-treated blood group, CDK6, NKG7, and LEF1 are markers of immune response and cell cycle regulation, indicating immune suppression. **(D)** In the paclitaxel and atezolizumab-treated blood group, SEC61B, CSF3R, and HLA-DRB1 reflect enhanced immune responses in macrophages, monocytes, and T cells.

On the other hand, in the only paclitaxel-treated blood group, genes such as *CDK6, NKG7*, and *LEF1,* which are involved in cell cycle regulation and immune response, were identified as advanced markers of immune response to paclitaxel treatment. These genes indicate cellular activity and immune suppression in response to treatment (Figure 6C). In contrast, in the blood group treated with combination therapy, paclitaxel and atezolizumab, genes such as *SEC61B*, *CSF3R* and *HLA-DRB1*, which refer to the immune responses of macrophages, monocytes and T cells, were identified (Figure 6D). These genes are involved in the active immune responses of the blood due to the combination therapy of paclitaxel and atezolizumab and indicate the positive effects of this combination therapy on immune responses in the blood (Supplementary Table 5).

Generally, while the tumor groups are biased towards markers of immune exhaustion and toxicity, such as *CD8A* and *GZMB*, the blood groups emphasize the enhancement of immune responses and cell proliferation, which is characterized by genes such as SEC61B and CDK6. These results identify key biomarkers that can be used to assess the response to only paclitaxel therapy and the effect of the combination therapy of atezolizumab.

### Machine Learning-Driven Prioritization of Response-Associated Biomarkers

To identify the most informative predictors of treatment response, we employed a Random Forest (RF) framework, a nonlinear classification approach optimized for high-dimensional transcriptomic data. To ensure the findings were technically sound and resistant to cell-level noise, the models were trained on per-cell expression matrices using scaled transcriptomic values (Seurat scale.data) restricted to a curated biomarker candidate list. Each cell was assigned the clinical response label of its originating patient. To ensure the stability of the prioritized features, we performed 100 bootstrap iterations, ranking genes by their selection frequency across the ensembles. For each of the four clinical subgroups (Tumor vs. PBMC; Paclitaxel vs. Combination), the top 20 genes with the highest cumulative selection counts were isolated (Figure 7, Supplementary Table 6).

**Figure 7:**
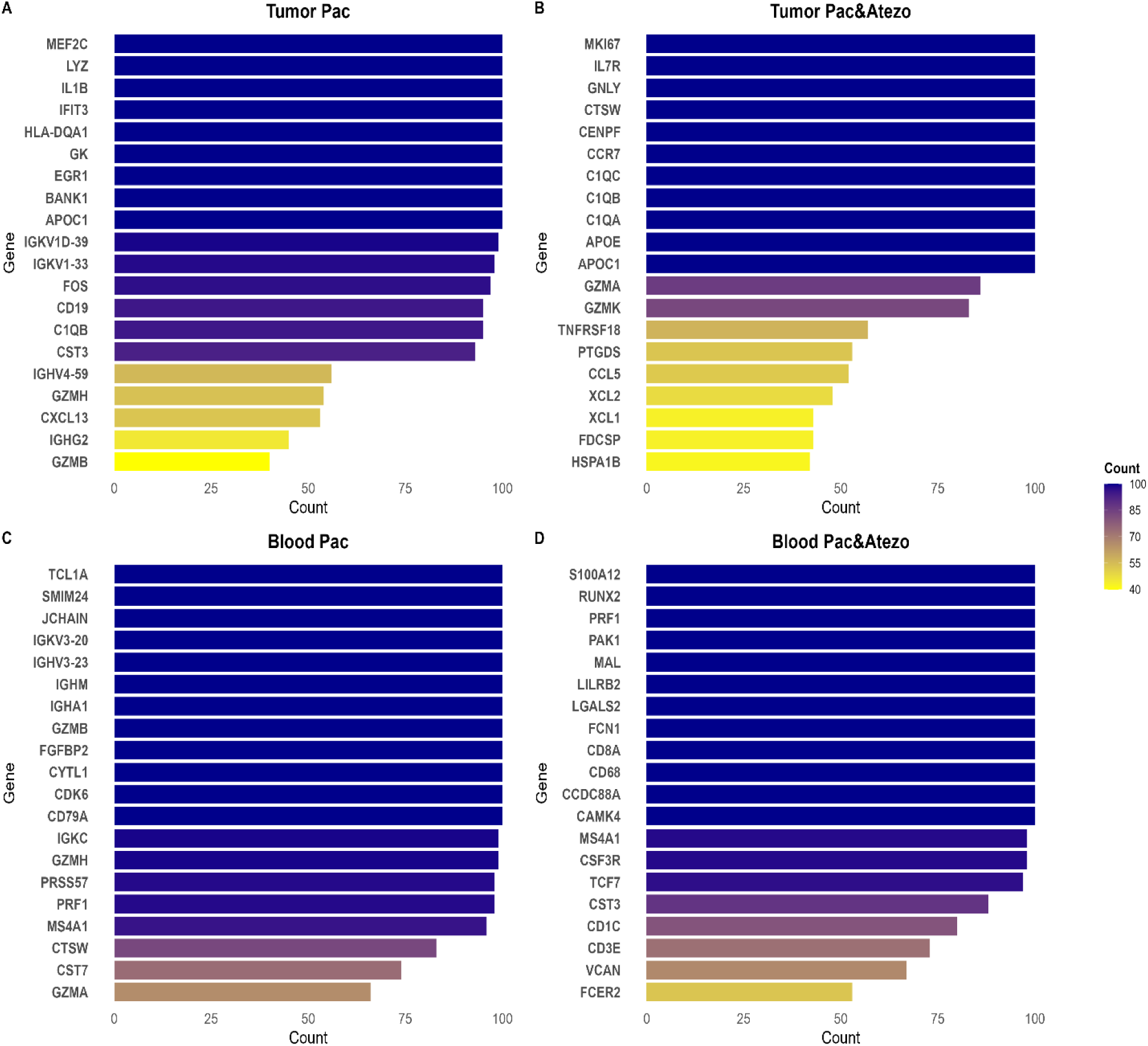
Top Genes Identified through Random Forest. This figure displays the top 20 genes identified using Random Forest analysis across four different treatment groups in TNBC patients. The genes are ranked based on their Count values, which reflect their importance in predicting treatment responses. The color gradient in each plot ranges from yellow (indicating lower values) to purple (indicating higher values), showing the intensity of each gene’s predictive significance. Count represents the number of 100 bootstrap iterations in which a gene was selected as an important Random Forest feature. **(A)** Tumor group treated with Paclitaxel. **(B)** Tumor group treated with Paclitaxel and Atezolizumab. **(C)** Blood group treated with Paclitaxel. **(D)** Blood group treated with Paclitaxel and Atezolizumab.

The resulting biomarker panels revealed a striking divergence between local and systemic compartments. In tumors treated with paclitaxel monotherapy, the predictive signature was dominated by inflammatory and stress-response markers, including *IL1B*, *IFIT3*, and *EGR1*, as well as myeloid-associated *LYZ* and *APOC1* (Figure 7A). Conversely, the combination therapy group shifted the predictive landscape toward a more integrated immune-proliferative state. Top predictors in this cohort included the complement components *C1QA/B/C*, the proliferation markers *MKI67* and *CENPF*, and the immune-trafficking receptor *CCR7* (Figure 7B).

Systemic readouts in the peripheral blood similarly distinguished the two regimens. Paclitaxel monotherapy predictors were heavily enriched for B-lineage and cytotoxic signatures, specifically immunoglobulin chains (*IGHA1/IGHM/IGKV*), *CD79A*, and the effector *GZMB* (Figure 7C). In the combination therapy arm, however, the model prioritized a broader myeloid and T-cell signaling network, featuring *S100A12*, *CD8A*, *PRF1*, and the regulatory transcripts *MAL* and *CAMK4* (Figure 7D).

This marked divergence underscores those therapeutic mechanisms, specifically the addition of PD-L1 blockade, which fundamentally rewire the transcriptional markers of response. Our findings are grounded in existing biological literature: *APOC1* is a recognized prognostic factor in TNBC (Yan et al., 2022), and the *C1Q* complex has been previously linked to favorable immune infiltration and patient survival (Mangogna et al., 2019). Furthermore, the emergence of cytotoxic effector markers like *GZMB* and *PRF1* in the circulation suggests that non-invasive, blood-based biopsies may serve as viable proxies for monitoring intra-tumoral immune engagement (Guan et al., 2024). Collectively, these subgroup-specific gene sets provide a high-fidelity panel for the development of personalized predictive signatures in TNBC.

### Evaluation of K-NN Classifier Performance

To evaluate whether the candidate biomarkers prioritized by Random Forest retained predictive utility in an independent modeling framework, we implemented a bootstrap-enhanced K-nearest neighbors (K-NN) classifier for each treatment subgroup. The primary objective was to quantify classification performance for treatment response using accuracy, sensitivity, specificity, and area under the receiver operating characteristic curve (AUC). To obtain stable performance estimates and reduce dependence on any single train/test split, we performed 100 bootstrap iterations per subgroup and report the mean ± standard deviation across iterations.

Across both tumor and peripheral blood/PBMC datasets, the K-NN classifier achieved consistently high discriminative performance using the selected gene panels (Figure 8, Supplementary Table 7). In tumor samples treated with paclitaxel alone, the model achieved accuracy = 0.988 ± 0.003 and AUC = 0.992 ± 0.005 (Figure 8A), indicating strong separation between complete responders and partial/non-responders in this subgroup. In tumor samples treated with paclitaxel plus atezolizumab, performance remained high (accuracy = 0.949 ± 0.015; AUC = 0.975 ± 0.011; Figure 8B), supporting robust classification under the combination regimen.

**Figure 8:**
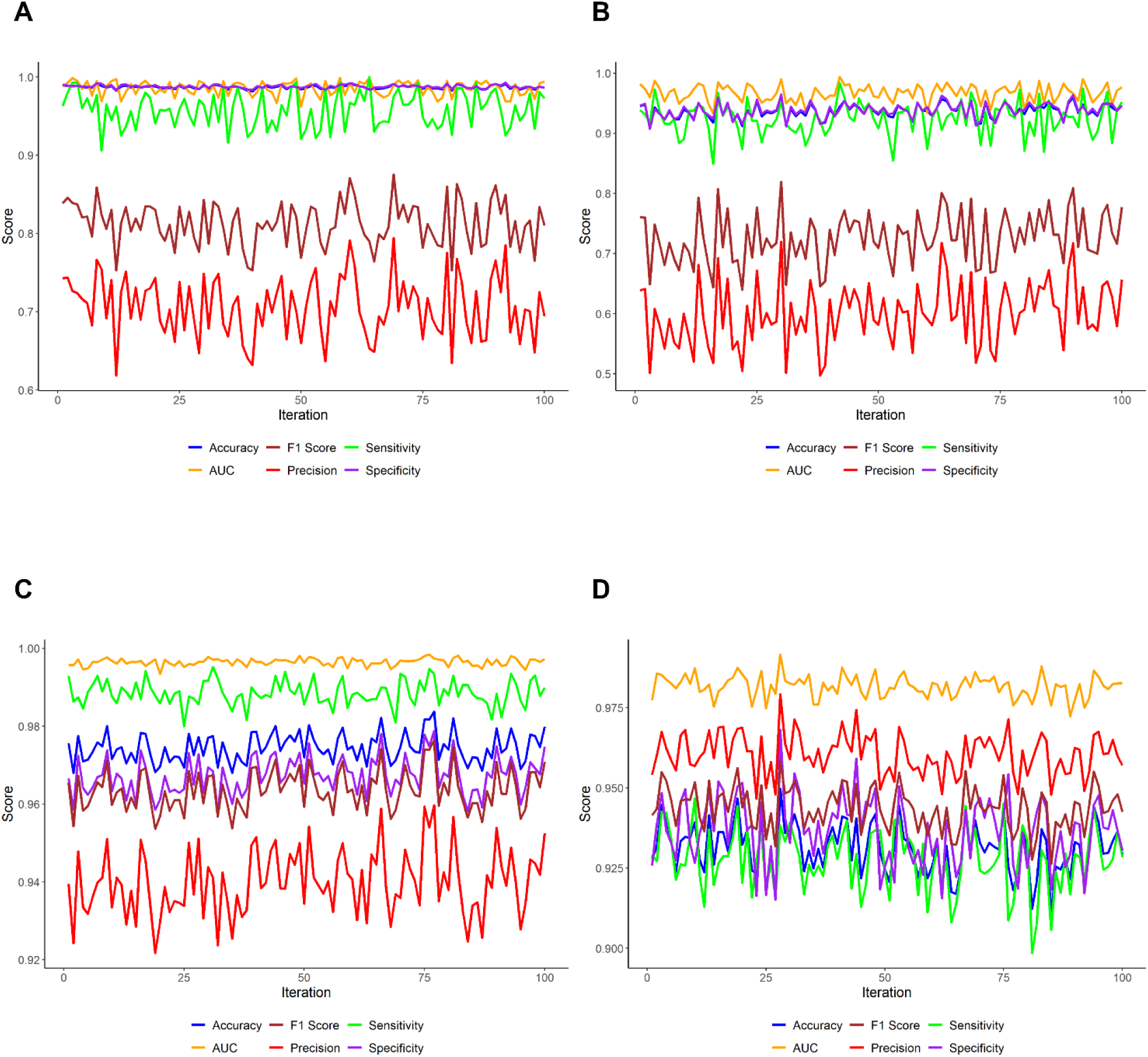
K-NN Classifier Performance Evaluation. It presents the performance of a K-NN classifier in predicting treatment response based on gene biomarkers identified through the Random Forest algorithm. The classifier’s accuracy, sensitivity, specificity, and AUC were evaluated across four different treatment groups. **(A)** This plot demonstrates the K-NN classifier’s exceptional performance on tumor samples treated with only paclitaxel. **(B)** This plot shows the model’s reliable performance in tumor samples treated with paclitaxel and atezolizumab. **(C)** The K-NN classifier performed even better in blood samples treated with paclitaxel. This demonstrates the potential of the model to predict treatment outcomes non-invasively, using blood-based samples, which is a promising development for liquid biopsies. **(D)** In blood samples treated with both paclitaxel and atezolizumab, the classifier showed slightly lower accuracy, but still demonstrated robust predictive power even under complex therapeutic conditions.

High performance was also observed in peripheral blood/PBMC samples. For blood/PBMC treated with paclitaxel alone, the classifier achieved accuracy = 0.974 ± 0.005 and AUC = 0.995 ± 0.002 (Figure 8C), indicating that circulating expression signatures can provide strong discriminatory signal in this setting. For blood/PBMC treated with paclitaxel plus atezolizumab, performance remained strong (accuracy = 0.931 ± 0.005; AUC = 0.985 ± 0.002; Figure 8D), consistent with stable predictive signal despite regimen-associated immune remodeling (Supplementary Table 7).

Collectively, these results indicate that the Random Forest–prioritized gene panels generalize to a distinct classifier and yield reproducible prediction performance across tumor and blood compartments and across both treatment regimens. The consistently high AUC values across all four subgroups support the utility of these biomarker sets as candidates for downstream validation and for development of response-prediction signatures in TNBC. Notably, the strong performance in blood/PBMC suggests the feasibility of minimally invasive, circulation-based monitoring strategies, although external validation in independent cohorts and harmonized sampling designs will be required to establish clinical generalizability.

### Interpretation and Selection of Top Genes Using LIME

To improve the interpretability of the response-prediction framework, we applied LIME (Local Interpretable Model-Agnostic Explanations) to quantify how individual genes influence K-NN predictions when restricting features to the top 20 genes prioritized by Random Forest. This analysis provides local, sample-level explanations that help identify which features most strongly push predictions toward the responder versus non-responder class within each treatment–sample subgroup (Figure 9).

**Figure 9.**
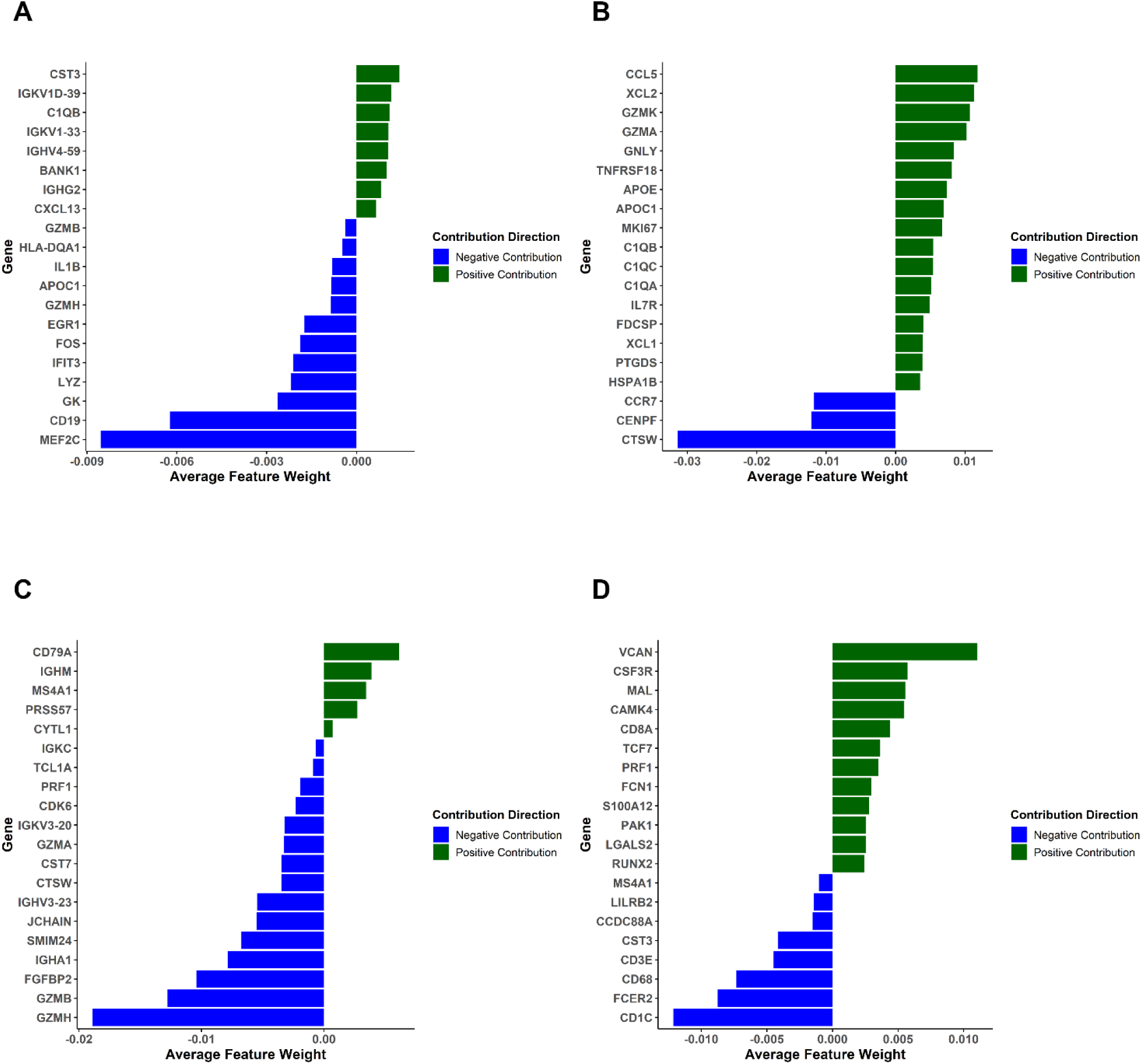
**A to D:** Each part presents an analysis of gene importance in predicting treatment response for different conditions. **(A)** Tumor samples treated with paclitaxel, the genes CST3 and IGKV1D-39 appear as key positive predictors of treatment sensitivity, with high expression linked to a greater likelihood of a positive treatment response, while, genes like MEF2C, GK, and APOC1 are associated with negative contribution, indicating resistance to paclitaxel. **(B)** Tumor samples treated with paclitaxel and atezolizumab, CCL5 and XCL2 (proliferation-related genes) show strong positive contributions, indicating that tumors with high proliferation rates respond better to the combination therapy. Negative predictors such as CTSW and CENPF are linked to immune evasion mechanisms that reduce therapeutic efficacy. **(C)** Blood samples treated with paclitaxel, Genes like CD79A, IGHM, and MS4A1 positively influence treatment response. **(D)** Blood samples treated with paclitaxel and atezolizumab, key immune-related genes such as VCAN, CD8A, and PRF1 show strong contributions, highlighting the role of T-cell mediated cytotoxicity in improving therapeutic response. Negative contributors like CD1C and FCER2 indicate the presence of suppressive immune mechanisms that might limit therapy effectiveness.

In tumor samples treated with paclitaxel alone, LIME highlighted *CST3* and *IGKV1D-39* as influential contributors to responder classification in the local explanation profiles, indicating that higher expression of these genes tended to increase the predicted probability of response in this subgroup (Figure 9A). Among the most prominent positive contributors, *EGR1* exerted the most substantial effect on responder prediction, consistent with prior work linking *EGR1* to breast cancer proliferation and drug sensitivity and implicating it in paclitaxel resistance mechanisms through its relationship with *YB-1* (Lasham et al., 2016). In addition, *EGR1* has been reported to promote ferroptosis and suppress aggressive phenotypes via *NRF2–HMOX1* signaling (Lin et al., 2024), supporting biological plausibility for its association with favorable classification. *IFIT3* also showed a consistent positive contribution, albeit with lower weight, aligning with its described roles in innate immune signaling and tumor biology (Wu et al., 2025). Conversely, in this subgroup, the strongest negative contributor was *MEF2C*, with *GK* and *APOC1* also showing negative contributions, indicating that higher expression of these features tended to shift predictions toward non-response (Figure 9A).

In tumor samples treated with paclitaxel plus atezolizumab, LIME explanations emphasized immune and cell-state programs as major drivers of classification. Chemokine genes such as *CCL5* and *XCL2* carried strong positive contributions, alongside cytotoxic/immune-associated genes (*GZMK, GNLY*) and complement components (*C1QA/B/C*) (Figure 9B). Notably, *MKI67* (KI-67), a canonical proliferation marker, also contributed positively to the combination subgroup, consistent with reports associating higher *MKI67* levels with chemotherapy responsiveness (Kim, K. I., et al., 2014).

Complement-associated programs have also been linked to anti-tumor immune activity in TNBC contexts (Yang et al., 2022), supporting the relevance of *C1QA/B/C* as interpretable contributors in this setting. In contrast, genes such as *CTSW* and *CENPF* showed pronounced negative contributions in the LIME profiles, indicating that higher expression of these features tended to shift model output toward non-response under combination therapy (Figure 9B). Together, these patterns suggest that under chemo-immunotherapy, classification is strongly influenced by variation in immune signaling and cell-state features, including cytotoxic activity and proliferative programs, rather than a single dominant transcript.

We also evaluated LIME explanations in peripheral blood/PBMC because blood-based signatures offer a plausible route to minimally invasive monitoring, including liquid biopsy strategies in TNBC (Mazzeo et al., 2024). In blood samples treated with paclitaxel alone, the strongest positive contributors to responder classification were *CD79A, IGHM, and MS4A1*, consistent with the subgroup’s prominent B-lineage features. In contrast, features including *TCL1A, JCHAIN,* and *CDK6* contributed negatively to the local explanation profiles, indicating association with non-response classification in this dataset. Additional negative contributors included *GZMH, FGFBP2*, and *IGKV3-20*, which also shifted predictions toward non-response when highly expressed (Figure 9C). Taken together, these results indicate that under paclitaxel monotherapy, the blood-based classifier is influenced by a balance of B-lineage programs and cytotoxic/effector-associated signatures, with distinct subsets contributing to opposite directions in the model’s local decision rules.

In blood/PBMC samples treated with paclitaxel plus atezolizumab, LIME highlighted a set of features consistent with cytotoxic T-cell activity and immune signaling. *CD8A* and *PRF1* emerged as top positive contributors, consistent with the interpretation that stronger cytotoxic effector programs support responder classification under combined chemo-immunotherapy (Figure 9D) and align with prior reports linking cytotoxic T-cell activity to improved outcomes in TNBC immunotherapy settings (Takabe et al., 2021). Additional influential features included *CAMK4*, which has been implicated in T-cell activation and immune transcriptional regulation (Koga et al., 2018), and *VCAN*, which may relate to extracellular matrix-associated programs and immune cell trafficking (Figure 7D). LIME also identified signaling and cytoskeletal-associated predictors, including *PAK1* and *CCDC88A* (Girdin), as contributors with opposite directions in the explanation profiles, consistent with prior work implicating PAK signaling in actin remodeling and invasion (Best et al., 2022) and Girdin in AKT-linked cytoskeletal integrity and migration (Enomoto et al., 2005). Conversely, *CD1C* and *FCER2* showed strong negative contributions, suggesting that higher expression of dendritic/B-cell immunoregulatory features may be associated with non-response classification in this subgroup (Figure 9D), consistent with published links between *CD1C^+^* dendritic subsets and immunoregulatory programs (Villani et al., 2017) and FCER2-related regulatory signaling in B-cell biology (Chan et al., 2014).

Collectively, LIME-based interpretation indicates that the genes driving predictions are both compartment- and regimen-dependent, with tumor predictions emphasizing chemokine/complement and proliferative programs under combination therapy, and blood predictions highlighting cytotoxic T-cell effector features under chemo-immunotherapy and B-lineage-associated programs under paclitaxel monotherapy. These LIME-selected genes refine the Random Forest feature lists by prioritizing features that consistently influence model decisions in biologically interpretable directions (Figure 9). Given the local nature of LIME and potential correlation among expression features, this candidate biomarkers should be validated in independent cohorts, ideally with patient-level partitioning to avoid information leakage and with orthogonal assays (e.g., targeted expression panels or protein-level measurements) to support translation into robust predictive signatures.

LIME explanations were generated using cell-level gene expression features (scaled Seurat expression values) and were applied only to held-out test cells after model training. For each subgroup, 100 test cells were randomly sampled for explanation. Gene-level LIME importance was summarized by aggregating feature contributions across the explained test cells, and the average (optionally absolute) LIME weight was reported per gene to highlight consistent directional contributors.

LIME was used solely as a post hoc interpretability tool and did not further filter or select features beyond the top-20 genes identified by Random Forest. Therefore, the final retained gene set per subgroup remained the Random Forest top-20 list, while LIME was used to interpret directionality and relative contribution. We note that assigning patient-level labels to single cells and splitting at the cell level may introduce optimistic estimates due to shared patient-specific signals. Future analyses will prioritize patient-level partitioning (or pseudo-bulk summaries) to fully mitigate potential information leakage.

## 4. Discussion

In this study, a comprehensive single-cell transcriptomic framework integrating differential gene expression, CNV analysis, co-expression network reconstruction, functional enrichment, and machine learning-based biomarker prioritization to uncover molecular determinants of treatment response in triple-negative breast cancer was applied. By comparing tumor- and blood-derived single cells from patients treated with paclitaxel alone or in combination with atezolizumab, high-resolution immune and tumor cell maps were generated that revealed distinct transcriptional adaptations under different therapeutic regimens.

Our findings provide several novel insights. First, the significant remodeling of the tumor microenvironment following treatment was identified by the clustering analysis. Treatment with paclitaxel alone activated cytotoxic T cells, exhausted *CD8⁺ T* cells, and macrophage-associated transcriptional programs, and correlated with prior reports indicating chemotherapy-induced immune infiltration in TNBC (Kalinski et al., 2025; Zhao et al., 2023). By contrast, the combination therapy with atezolizumab was associated with augmented B-cell expansion, plasmablast differentiation, and a lack of activated T cells, showing a potential shift in immune compartmentalization toward humoral immunity. These results align with prior clinical observations that checkpoint blockade enhances B-cell–mediated responses in breast cancer (Flippot et al., 2024). In addition, differential expression and pathway enrichment across tumor and blood/PBMC compartments delineated both shared immune-associated themes and regimen-specific transcriptional programs. The tumor compartment exhibited immune-centric signatures in both arms, with broader cytokine/chemokine-related pathway enrichment in the combination therapy group, whereas peripheral blood/PBMC profiles highlighted treatment-associated remodeling involving innate/myeloid markers, mast cell-associated transcripts, and humoral/T-helper differentiation programs.

Collectively, the studies on differential expression and pathway analyses indicated that combination therapy could mobilize and stimulate more diverse and complex signaling programs than any type of monotherapy. For example, pathways enriched in the paclitaxel-alone-treated group include leukocyte-mediated immunity and lymphocyte chemotaxis; treatment with paclitaxel in combination with atezolizumab activated pathways such as chemokine signaling and viral proteins interacting with cytokines, thus suggesting wider immune stimulation. Thus, it provides support to the concept that immunotherapy can change the immune signature induced by chemotherapy, with a more apt effect of such changes on long-term activity against tumors (Ademuyiwa et al., 2022).

On the other hand, changes in immune-related genes, such as *IL7R, CD6,* and *TNFAIP3*, especially in T cell populations, were analyzed ing CNV analysis. Altered expression of *IL7R, CD6,* and *TNFAIP3* post-treatment makes them attractive candidates for predictive biomarkers of immunotherapy benefit. *IL7R* is a key gene for T-cell development and homeostasis, and its alteration leads to dysfunctional anti-tumor immunity and resistance to therapy (Lin, 2017). The co-stimulation receptor CD6, which is expressed in T cells, has been reported to facilitate immune synapse formation and T-cell activation, thereby enhancing T cell responses during immune checkpoint blockade (Gonçalves, 2018). In cytokine signaling, *TNFAIP3* (also termed *A20*) serves as a crucial negative regulator of the *NF-kB* pathway, mutations in this gene have been linked to immune escape and responses to immunotherapy (Catrysse, L. et al., 2014).

Last but not least, these genes were integrated into WGCNA and found to cluster into treatment-specific co-expression modules, strengthening their biological plausibility as major players in the immunological response to therapy. Together, these observations combined lend strong support to the importance of CNV profiling in network approaches for consistently robust therapeutic response biomarkers.

Machine learning took this a step further, refining the biomarker landscape by prioritizing the genes most predictive of treatment response. Just as *IGKC* and *IGHG1*, immune globulin-related genes, and other macrophage-associated genes, *APOE* and *CST3*, were repeatedly identified across statistical and computational analyses, it becomes increasingly certain that connections across independent analytic layers could enhance their clinical utility as predictive biomarkers. This study concludes that combining single-cell multi-omics with network and machine learning approaches can reveal intricate details treatment responses in TNBC. Discovery of shared and therapy-specific biomarkers provides further scope for precision oncology, enabling patients to be classified based on molecular predictors of response to chemotherapy or chemo-immunotherapy.

However, several limitations must be acknowledged. Firstly, the single-cell RNA sequencing (scRNA-seq) datasets utilized in this study are constrained by limited sample sizes and heterogeneity. Secondly, the functional validation of the identified biomarkers was beyond the scope of this investigation but represents a crucial avenue for future research to elucidate their causal roles in treatment response. Although our computational pipeline yielded robust biomarker candidates, the validation in independent cohorts was limited by the availability of datasets. At the time of analysis, no publicly accessible datasets with comparable experimental design and clinical parameters (pre- and post-treatment samples under paclitaxel and paclitaxel combined with atezolizumab regimens) were available. Given the enhancement of biomarker prioritization through machine learning, the inclusion of independent validation cohorts is essential to ensure the generalizability of these findings. With this, the research provides a well-structured framework for identifying biomarker candidates that are likely to predict how triple-negative breast cancer responds to treatment. Although only a few biomarkers and pathways have received attention, they appear promising for patient stratification and for optimizing combination regimens in treatment, particularly in pathologies dependent on immune modulation. Future prospective studies will be essential for translating these findings into personalized therapy for TNBC by incorporating scRNA-seq profiling with clinical outcome data.

## References

1. Xu K, Wang R, Xie H, Hu L, Wang C, Xu J, et al. (2021). Single-cell RNA sequencing reveals cell heterogeneity and transcriptome profile of breast cancer lymph node metastasis. Oncogenesis. 10(10):66. doi:10.1038/s41389-021-00355-6

2. Cai Y, Dai F, Ye Y, Qian J. (2025). The global burden of breast cancer among women of reproductive age: a comprehensive analysis. Sci Rep. 15(1):9347. doi:10.1038/s41598-025-93883-9

3. Jain P, Aggarwal S, Adam S, Imam M. (2024). Parametric optimization and comparative study of machine learning and deep learning algorithms for breast cancer diagnosis. Breast Dis. 43(1):257–270. doi:10.3233/BD-240018

4. Cummings-John C, Bah AJ, Smalle IO, Challe CC, Duduyemi B, Ogundiran T. (2025). The patterns of presentation, management and outcomes of breast cancer patients at a tertiary health facility in Sierra Leone. BMC Cancer. 25(1):70. doi:10.1186/s12885-025-13429-0

5. Baba AI, Catoi C. (2007). Chapter 3. Tumor Cell Morphology. In: Buchares, R.O., Ed., Comparative Oncology. The Publishing House of the Romanian Academy, Bucharest.

6. Turashvili G, Brogi E. (2017). Tumor Heterogeneity in Breast Cancer. Frontiers in medicine, 4, 227. doi:10.3389/fmed.2017.00227

7. Livesey BJ, Marsh JA. (2022). The properties of human disease mutations at protein interfaces. PLoS Comput Biol. 18(2):e1009858. doi:10.1371/journal.pcbi.1009858

8. Hausser J, Szekely P, Bar N, et al. (2019). Tumor diversity and the trade-off between universal cancer tasks. Nat Commun. 10, 5423. doi:10.1038/s41467-019-13195-1

9. Ahmed M, Kim DR. (2024). Life history dynamics of evolving tumors: insights into task specialization, trade-offs, and tumor heterogeneity. Cancer Cell Int. 24(1):364. doi:10.1186/s12935-024-03538-4

10. Garg P, Malhotra J, Kulkarni P, Horne D, Salgia R, Singhal SS. (2024). Emerging Therapeutic Strategies to Overcome Drug Resistance in Cancer Cells. Cancers (Basel). 16(13):2478. doi:10.3390/cancers16132478

11. Wang Y, Chen Q, Shao H, Zhang R, Shen H. (2024). Generating bulk RNA-Seq gene expression data based on generative deep learning models and utilizing it for data augmentation. Comput Biol Med. 169:107828. doi:10.1016/j.compbiomed.2023.107828

12. Zhang, Y., Wang, D., Peng, M., Tang, L., Ouyang, J., Xiong, F., Guo, C., Tang, Y., Zhou, Y., Liao, Q., Wu, X., Wang, H., Yu, J., Li, Y., Li, X., Li, G., Zeng, Z., Tan, Y., & Xiong, W. (2021). Single-cell RNA sequencing in cancer research. Journal of experimental & clinical cancer research : CR, 40(1), 81. doi:10.1186/s13046-021-01874-1

13. Sun G, Li Z, Rong D, Zhang H, Shi X, Yang W, Zheng W, Sun G, Wu F, Cao H, Tang W, Sun Y. (2021). Single-cell RNA sequencing in cancer: Applications, advances, and emerging challenges. Mol Ther Oncolytics. 21:183–206. doi:10.1016/j.omto.2021.04.001

14. Cruz, J. A., & Wishart, D. S. (2007). Applications of machine learning in cancer prediction and prognosis. Cancer informatics, 2, 59–77.

15. Zhang, Y., Chen, H., Mo, H., Hu, X., Gao, R., Zhao, Y., Liu, B., Niu, L., Sun, X., Yu, X., Wang, Y., Chang, Q., Gong, T., Guan, X., Hu, T., Qian, T., Xu, B., Ma, F., Zhang, Z., & Liu, Z. (2021). Single-cell analyses reveal key immune cell subsets associated with response to PD-L1 blockade in triple-negative breast cancer. Cancer cell, 39(12), 1578–1593.e8. 10.1016/j.ccell.2021.09.010

16. Pal, B., Chen, Y., Vaillant, F., Capaldo, B. D., Joyce, R., Song, X., Bryant, V. L., Penington, J. S., Di Stefano, L., Tubau Ribera, N., Wilcox, S., Mann, G. B., kConFab, Papenfuss, A. T., Lindeman, G. J., Smyth, G. K., & Visvader, J. E. (2021). A single-cell RNA expression atlas of normal, preneoplastic and tumorigenic states in the human breast. The EMBO journal, 40(11), e107333. doi10.15252/embj.2020107333

17. Gambardella G, et al. (2022). A single-cell analysis of breast cancer cell lines to study tumour heterogeneity and drug response. Nature communications, 13(1), 1714.

18. Chen, Y., Pal, B., Lindeman, G.J. et al. R code and downstream analysis objects for the scRNA-seq atlas of normal and tumorigenic human breast tissue. Sci Data 9, 96 (2022). doi10.1038/s41597-022-01236-2

19. Hao, Y., Hao, S., Andersen-Nissen, E., Mauck, W. M., 3rd, Zheng, S., Butler, A., Lee, M. J., Wilk, A. J., Darby, C., Zager, M., Hoffman, P., Stoeckius, M., Papalexi, E., Mimitou, E. P., Jain, J., Srivastava, A., Stuart, T., Fleming, L. M., Yeung, B., Rogers, A. J., Satija, R. (2021). Integrated analysis of multimodal single-cell data. Cell, 184(13), 3573–3587.e29. doi:10.1016/j.cell.2021.04.048

20. Hafemeister, C., & Satija, R. (2019). Normalization and variance stabilization of single-cell RNA-seq data using regularized negative binomial regression. Genome biology, 20(1), 296. doi:10.1186/s13059-019-1874-1

21. McInnes L, Healy J, Saul N, Großberger L. (2018). UMAP: Uniform Manifold Approximation and Projection. Journal of Open Source Software. 2020;5(53):2185. doi:10.21105/joss.00861

22. Zhang, X., Lan, Y., Xu, J., Quan, F., Zhao, E., Deng, C., Luo, T., Xu, L., Liao, G., Yan, M., Ping, Y., Li, F., Shi, A., Bai, J., Zhao, T., Li, X., & Xiao, Y. (2019). CellMarker: a manually curated resource of cell markers in human and mouse. Nucleic acids research, 47(D1), D721–D728. doi:10.1093/nar/gky900

23. Patel AP, Tirosh I, Trombetta JJ, et al. (2014). Single-cell RNA-seq highlights intratumoral heterogeneity in primary glioblastoma. Science. 344(6190):1396–1401. doi:10.1126/science.1254257

24. Wilcoxon F. (1946). Individual comparisons of grouped data by ranking methods. Journal of economic entomology, 39, 269. doi:10.1093/jee/39.2.269

25. Cunningham, F., Allen, J. E., Allen, J., Alvarez-Jarreta, J., Amode, M. R., Armean, I. M., Austine-Orimoloye, O., Azov, A. G., Barnes, I., Bennett, R., Berry, A., Bhai, J., Bignell, A., Billis, K., Boddu, S., Brooks, L., Charkhchi, M., Cummins, C., Da Rin Fioretto, L., Davidson, C., … Flicek, P. (2022). Ensembl 2022. Nucleic acids research, 50(D1), D988–D995. doi:10.1093/nar/gkab1049

26. Nguyen, P., & Zeng, E. (2025). A Protocol for Weighted Gene Co-expression Network Analysis With Module Preservation and Functional Enrichment Analysis for Tumor and Normal Transcriptomic Data. Bio-protocol, 15(18), e5447. doi10.21769/BioProtoc.5447

27. Horvath, S. (2011). “Weighted network analysis: applications in genomics and systems biology.” Springer Science & Business Media.

28. Langfelder, P., & Horvath, S. (2008). WGCNA: an R package for weighted correlation network analysis. BMC bioinformatics, 9, 559. doi:10.1186/1471-2105-9-559

29. Barabasi, A. L., & Albert, R. (1999). Emergence of scaling in random networks. Science (New York, N.Y.), 286(5439), 509–512. doi:10.1126/science.286.5439.509

30. Gene Ontology Consortium (2021). The Gene Ontology resource: enriching a GOld mine. Nucleic acids research, 49(D1), D325–D334. doi10.1093/nar/gkaa1113

31. Kanehisa, M., Furumichi, M., Sato, Y., Ishiguro-Watanabe, M., & Tanabe, M. (2021). KEGG: integrating viruses and cellular organisms. Nucleic acids research, 49(D1), D545–D551. doi:10.1093/nar/gkaa970

32. Yu, G., Wang, L. G., Han, Y., & He, Q. Y. (2012). clusterProfiler: an R package for comparing biological themes among gene clusters. Omics: a journal of integrative biology, 16(5), 284–287. doi:10.1089/omi.2011.0118

33. Huber, W., Carey, V. J., Gentleman, R., Anders, S., Carlson, M., Carvalho, B. S., … & Smyth, G. K. (2015). Orchestrating high-throughput genomic analysis with Bioconductor. Nature Methods, 12(2), 115–121. doi10.1038/nmeth.3252

34. Moorthy, K., & Mohamad, M. S. (2011). Random forest for gene selection and microarray data classification. Bioinformation, 7(3), 142–146. doi10.6026/97320630007142

35. Efron, B., & Tibshirani, R. (1997). Improvements on Cross-Validation: The 632+ Bootstrap Method. Journal of the American Statistical Association, 92(438), 548–560. doi10.1080/01621459.1997.10474007

36. Díaz-Uriarte, R., & Alvarez de Andrés, S. (2006). Gene selection and classification of microarray data using random forest. BMC bioinformatics, 7, 3. 10.1186/1471-2105-7-3

37. N. V. Chawla, K. W. Bowyer, L. O. Hall, W. P. Kegelmeyer. (2002). SMOTE: Synthetic Minority Over-sampling Technique. Journal of Artificial Intelligence Research 16 (2002) 321–357. doi:10.1613/jair.953

38. Yan, et al. (2022). Apolipoprotein C1 (APOC1), A Candidate Diagnostic Serum Biomarker for Breast Cancer Identified by Serum Proteomics Study. Critical reviews in eukaryotic gene expression, 32(4), 1–9. doi:10.1615/CritRevEukaryotGeneExpr.2021040967

39. Mangogna, et al. (2019). Is the Complement Protein C1q a Pro- or Anti-tumorigenic Factor? Frontiers in immunology, 10, 865. doi:10.3389/fimmu.2019.00865

40. Voloshin, et al. (2015). Blocking IL1β Pathway Following Paclitaxel Chemotherapy Slightly Inhibits Primary Tumor Growth but Promotes Spontaneous Metastasis. Molecular cancer therapeutics, 14(6), 1385–1394. doi:10.1158/1535-7163.MCT-14-0969

41. Wang, et al. (2020). Inflammatory IFIT3 renders chemotherapy resistance by regulating post-translational modification of VDAC2 in pancreatic cancer. Theranostics, 10(16), 7178–7192. doi:10.7150/thno.43093

42. Davey, et al. (2021). Ki-67 as a Prognostic Biomarker in Invasive Breast Cancer. Cancers, 13(17), 4455. doi:10.3390/cancers13174455

43. Tao, et al. (2021). The roles of the cell division cycle-associated gene family in hepatocellular carcinoma. Journal of gastrointestinal oncology, 12(2), 781–794. doi:10.21037/jgo-21-110

44. Hay, et al. (2022). Granzymes: The Molecular Executors of Immune-Mediated Cytotoxicity. International journal of molecular sciences, 23(3), 1833. doi:10.3390/ijms23031833. Guan et al. Perforin 1 in Cancer: Mechanisms, Therapy, and Outlook. Biomolecules, 14(8), 910 (2024). doi:10.3390/biom14080910

45. Guan, X., Guo, H., Guo, Y., Han, Q., Li, Z., & Zhang, C. (2024). Perforin 1 in Cancer: Mechanisms, Therapy, and Outlook. Biomolecules, 14(8), 910. doi10.3390/biom14080910

46. Lasham, et al. (2016). A novel EGR-1 dependent mechanism for YB-1 modulation of paclitaxel response in a triple negative breast cancer cell line. International journal of cancer, 139(5), 1157–1170. doi:10.1002/ijc.30137

47. Lin, et al. (2024). EGR1 Promotes Erastin-induced Ferroptosis Through Activating Nrf2-HMOX1 Signaling Pathway in Breast Cancer Cells. Journal of Cancer, 15(14), 4577–459. doi:10.7150/jca.95328

48. Wu, et al. (2025). IFIT3: a crucial mediator in innate immunity and tumor progression with therapeutic implications. Frontiers in immunology, 16, 1515718. doi:10.3389/fimmu.2025.1515718

49. Kim, K. I., Lee, K. H., Kim, T. R., Chun, Y. S., Lee, T. H., & Park, H. K. (2014). Ki-67 as a predictor of response to neoadjuvant chemotherapy in breast cancer patients. Journal of breast cancer, 17(1), 40–46. doi:10.4048/jbc.2014.17.1.40

50. Yang, et al. (2022). Prognostic and immune-related value of complement C1Q (C1QA, C1QB, and C1QC) in skin cutaneous melanoma. Frontiers in genetics, 13, 940306. doi:10.3389/fgene.2022.940306

51. Mazzeo, et al. (2024). Liquid biopsy in triple-negative breast cancer: unlocking the potential of precision oncology. ESMO open, 9(10), 103700 (2024). doi:10.1016/j.esmoop.103700

52. Takabe, et al. (2021). Immune cytolytic activity is associated with reduced intra-tumoral genetic heterogeneity and with better clinical outcomes in triple negative breast cancer. American journal of cancer research, 11(7), 3628–3644.

53. Koga, et al. (2018). The role of CaMK4 in immune responses. Modern rheumatology, 28(2), 211–214. doi:10.1080/14397595.2017.1413964

54. Best, et al. (2022). PAK-dependent regulation of actin dynamics in breast cancer cells. The international journal of biochemistry & cell biology, 146, 106207. doi:10.1016/j.biocel.2022.106207

55. Enomoto, et al. (2005). Akt/PKB regulates actin organization and cell motility via Girdin/APE. Developmental cell, 9(3), 389–402. doi:10.1016/j.devcel.2005.08.001

56. Villani, et al. (2017). Single-cell RNA-seq reveals new types of human blood dendritic cells, monocytes, and progenitors. Science (New York, N.Y.), 356(6335), eaah4573. doi:10.1126/science.aah4573

57. Chan, et al. (2014). FCER2 (CD23) asthma-related single nucleotide polymorphisms yields increased IgE binding and Egr-1 expression in human B cells. American journal of respiratory cell and molecular biology, 50(2), 263–269. doi:10.1165/rcmb.2013-0112OC

58. Kalinski P, Kokolus KM, Gandhi S. (2025). Paclitaxel, interferons and functional reprogramming of tumor-associated macrophages in optimized chemo-immunotherapy. Journal for immunotherapy of cancer, 13(5), e010960. doi:10.1136/jitc-2024-010960

59. Zhao H, Lin Z, Zhang Y, Liu J, Chen Q. (2023). Investigating the Heterogeneity of Immune Cells in Triple-Negative Breast Cancer at the Single-Cell Level before and after Paclitaxel Chemotherapy. International Journal of Molecular Sciences, 24(18), 14188. doi:10.3390/ijms241814188

60. Flippot R, Teixeira M, Rey-Cardenas M, Carril-Ajuria L, Rainho L, Naoun N, Jouniaux JM, Boselli L, Naigeon M, Danlos FX, Escudier B, Scoazec JY, Cassard L, Albiges L, Chaput N. (2024). B cells and the coordination of immune checkpoint inhibitor response in patients with solid tumors. Journal for immunotherapy of cancer, 12(4), e008636. doi:10.1136/jitc-2023-008636

61. Ademuyiwa FO, Gao F, Street CR, Chen I, Northfelt DW, Wesolowski R, Arora M, Brufsky A, Dees EC, Santa-Maria CA, Connolly RM, Force J, Moreno-Aspitia A, Herndon JM, Carmody M, Davies SR, Larson S, Pfaff KL, Jones SM, Weirather JL, et al. (2022). A randomized phase 2 study of neoadjuvant carboplatin and paclitaxel with or without atezolizumab in triple negative breast cancer (TNBC) - NCI 10013. NPJ breast cancer, 8(1), 134. doi:10.1038/s41523-022-00500-3

62. Lin JZ. (2017). The role of IL-7 in Immunity and Cancer. Anticancer research, 37(3), 963–967. doi:10.21873/anticanres.11405

63. Gonçalves CM. (2018). CD6, a Rheostat-Type Signalosome That Tunes T Cell Activation. Frontiers in immunology, 9, 2994. doi:10.3389/fimmu.2018.0299

64. Catrysse L, et al. (2014). A20 in inflammation and autoimmunity. Trends in immunology, 35(1), 22–31. doi:10.1016/j.it.2013.10.005

